# Systematic mapping of small nucleolar RNA targets in human cells

**DOI:** 10.1101/2021.07.22.451324

**Authors:** Hywel Dunn-Davies, Tatiana Dudnakova, Jean-Louis Langhendries, Nicholas Watkins, Denis L.J. Lafontaine, David Tollervey

## Abstract

Altered expression of box C/D small nucleolar RNAs (snoRNAs) is implicated in human diseases, including cancer. Box C/D snoRNAs canonically direct site-specific, 2’-*O*-methylation but the extent to which they participate in other functions remains unclear. To identify RNA targets of box C/D snoRNAs in human cells, we applied two techniques based on UV crosslinking, proximity ligation and sequencing of RNA hybrids (CLASH and FLASH). These identified hundreds of novel snoRNA interactions with rRNA, snoRNAs and mRNAs. We developed an informatic pipeline to rigorously call interactions predicted to direct methylation. Multiple snoRNA-rRNA interactions identified were not predicted to direct RNA methylation. These potentially modulate methylation efficiency and/or contribute to folding dynamics. snoRNA-mRNA hybrids included 1,300 interactions between 117 snoRNA families and 940 mRNAs. Human U3 is substantially more abundant than other snoRNAs and represented about 50% of snoRNA-mRNA hybrids. The distribution of U3 interactions across mRNAs also differed from other snoRNAs. Following U3 depletion, mRNAs showing altered abundance were strongly enriched for U3 CLASH targets. Most human snoRNAs are excised from pre-mRNA introns. Enrichment for snoRNA association with branch point regions of introns that contain snoRNA genes was common, suggesting widespread regulation of snoRNA maturation.

## INTRODUCTION

The small nucleolar RNAs (snoRNAs) are a class of abundant, small stable RNAs, most of which act as guides for site-specific RNA modification. Most members of the box C/D class of snoRNAs select sites of ribose 2’-*O*-methylation by extended regions of perfect complementarity with target sites (≥12 bp), in which the nucleotide to be modified is placed exactly 5 bp from the conserved box D or box D’ motifs within the snoRNA (reviewed in (Tollervey and Kiss, 1997; Watkins and Bohnsack, 2012)). The box C/D snoRNAs associate with a group of four common proteins, NOP56, NOP58, 15.5K and the methyltransferase Fibrillarin. The snoRNAs have a partially symmetrical structure, in which stem structures bring together the highly conserved, terminal box C (RUGAUGA, R = A or G) and box D (CUGA) sequences and the related, but less conserved, internal box C’ and box D’ elements. These stem structures include a K-turn structural motif that is bound by the small 15.5K protein. *In vitro* structural analysis indicated that the box C/D stem is also bound by NOP58, while the box C’/D’ stem is bound by the homologous NOP56 protein. Each region is bound by a copy of NOP1, so the regions flanking either box D, box D’ or both can function as methylation guides. In human cells, snoRNA-directed methylation sites show variable stoichiometry, indicating regulation that is likely to be functionally important (reviewed in (Henras et al., 2017; Sloan et al., 2017)).

The strict requirement for a long region of perfect complementarity that extends to box D/D’ for guide function implies that strong snoRNA base pairing could occur without eliciting target RNA methylation. Indeed, a small number of box C/D snoRNAs have essential functions in ribosome synthesis that require snoRNA/pre-rRNA base pairing without associated rRNA methylation. In vertebrates, these snoRNAs include U3, U14 and U8 (reviewed in (Watkins and Bohnsack, 2012)). In addition, pre-rRNA base-pairing by U13 snoRNA directs formation of *N*^4^-acetyl cytidine (ac4C) by NAT10, rather than methylation (Sharma et al., 2015; Sharma et al., 2017b).

Other acetylation targets are not known, but human snoRNAs can direct methylation of small RNAs, including spliceosomal small nuclear RNAs (snRNAs) and other snoRNAs (Bouchard-Bourelle et al., 2019; Bouchard-Bourelle et al., 2020; Dupuis-Sandoval et al., 2015). A number of human snoRNAs have been reported to bind mRNAs, with effects on pre-mRNA splicing and 3’ end formation (Falaleeva et al., 2016; Huang et al., 2017). Human snoRNAs show tissue-specific expression patterns, with altered expression linked to disease and tumorigenesis (Bergeron et al., 2021; Bratkovič et al., 2020; Fafard-Couture et al., 2021; Siprashvili et al., 2016; Su et al., 2013; Thorenoor and Slaby, 2015; Xu et al., 2014; Zhou et al., 2017). The imprinted, brain-specific snoRNAs snoRD115 and snoRD116 are implicated in the neurological disease Prader-Willi syndrome [see (Bieth et al., 2015; Burnett et al., 2017; Falaleeva et al., 2016; Falaleeva et al., 2015; Rozhdestvensky et al., 2016) reviewed in (Cavaillé, 2017)]. Mutations in U8/snoRD118 cause the neurological disease leukoencephalopathy with calcification and cysts in humans (Frenk et al., 2014) and a Zebrafish model (Badrock et al., 2020). In addition, snoRNAs can apparently function by direct protein binding: Loss of specific snoRNAs reduced levels of the GTP-bound, active form of K-Ras with consequent hyperactivation of the Ras-ERK1/ERK2 signaling pathway (Siprashvili et al., 2016). snoRNAs were also implicated in activation of the immune regulator Protein Kinase RNA-activated (PKR) under conditions of metabolic stress (Youssef et al., 2015).

Bioinformatics approaches have been used to predict snoRNA binding sites in several systems, particularly where this is associated with methylation (Jorjani et al., 2016; Lowe and Eddy, 1999; Lu et al., 2016; Omer et al., 2000). In addition, a number of recent reports have described methods for the identification of RNA-RNA interactions through proximity ligation followed by sequencing of the products of reverse transcription and PCR amplification (RT-PCR) (Bharathavikru et al., 2017; Dudnakova et al., 2018; Kudla et al., 2011; Sharma et al., 2016; Sugimoto et al., 2015). The <u>c</u>rosslinking <u>a</u>nd <u>s</u>equencing of <u>h</u>ybrids (CLASH) approach used stringent tandem affinity purification including denaturing conditions to recover RNA-protein and RNA-RNA interactions involving yeast snoRNAs (Dudnakova et al., 2018; Kudla et al., 2011) and human miRNAs (Helwak et al., 2013; Helwak and Tollervey, 2014). A related approach, formaldehyde-assisted crosslinking <u>a</u>nd <u>s</u>equencing of <u>h</u>ybrids (FLASH) combines immunoprecipitation with mild chemical crosslinking, which stabilizes protein complexes allowing denaturing wash conditions (Bharathavikru et al., 2017). Here we report the use of CLASH and FLASH to systematically map the interactions between box C/D snoRNAs and the human transcriptome.

## MATERIALS AND METHODS

### Human cell culture and UV crosslinking

Human embryonic kidney HEK cells were gown to 80% confluency in DMEM, 10%FBS medium and were UV crosslinked on ice with λ = 254 nm in Stratalinker 1800 (Stratagene), at 400 mJ/cm2. U3 was depleted from HEK cells by use of a chimeric antisense oligonucleotide, as described (Langhendries et al., 2016).

### HEK Cell Lysis

HEK cells were lysed by addition of ice-cold TM150 buffer (20 mM Tris-HCl pH 7.4, 150 mM NaCl, 0.4% NP-40, 2mM MgCl2, 1 mM DTT, protease inhibitors (Roche, complete, EDTA-free), RNAse Inhybitor (Promega)). 10u of RQ1DNAse (Promega) were added, the samples were mixed by pipetting and incubated for 10 min at room temperature to break genomic DNA and to ease the extraction of nuclear Fibrillarin binding complexes. Lysates were centrifuged in Eppendorf mini centrifuge at 14000rpm and 4C for 10 min and supernatant was collected.

### First affinity purification

In human HEK CLASH beads conjugated with anti-Flag M2 (Sigma), in FLASH Protein A agarose conjugated with IgG and anti-FBL AB were used. Cell lysates were incubated with beads for 60 min at 4°C. Supernatant was discarded and the recovered beads were washed twice with PBS-WB buffer (PBS, plus 150mM NaCl, 2 mM MgCl2, 0.4% NP-40), and once in 1X PBS, 2mM MgCl2.

### RNase treatment and formaldehyde crosslinking

RNP complexes bound to the beads were treated with 0.5 unit RNaseA+T1 mix (RNace-IT, Stratagene) in 100 μl PBS,2mM MgCl2 buffer for 10 min at 20°C. In CLASH 900 μl GDB denaturing buffer (6M GuCl2, 150mM NaCl, 20mM Tris pH=7.4) was added to the beads with RNase and mixed thoroughly. Supernatant with denatured complexes was removed and added to Ni beads (Gibco) washed in GDB, subsequent binding carried out for 1h at 4C. In FLASH to remove indirect RNA and protein binding from the complexes the beads were washed twice with PBS-WB buffer (PBS, +150mM NaCl, 2 mM MgCl2, 0.4% NP-40), twice with HS-PBS-WB (PBS, 0.3 M NaCl, 2 mM MgCl2, 0.4% NP-40) and once in 1xPBS. The complexes were cross linked on beads in 0.2% formaldehyde in PBS for 3 min, then formaldehyde was quenched by addition of glycine to 0.2M and Tris-HCl pH=8 to 0.1M and incubation for 5 min. Crosslinked complexes were subjected to 4x denaturing washes in UB (20 mM Tris pH=7.4, 8M UREA, 0.3M NaCl, 0.4% NP-40) and additional incubation in UB for 30 min at 4C to remove nonspecific interactions. Subsequent steps were identical for FLASH and CLASH.

### RNA end modification

In CLASH Ni beads were washed twice with GDB buffer, twice with TM150, twice with PNK buffer (50 mM Tris-HCl pH 7.5, 10 mM MgCl2, 0.5% NP-40, 50 mM NaCl). In FLASH beads with bound RNA-protein complexes were washed 4x with PNK buffer. To remove unwanted 3’phosphate groups from bound RNA fragments in both CLASH and FLASH the complexes were treated with TSAP phosphatase (Promega) using provided buffer for 40 min at room temperature. To inactivate the enzyme the beads were washed twice with UB (FLASH) or GDB (CLASH) and 4x with PNK buffer.

Then the 5’ phosphorylation and radioactive labelling of RNA was carried out. The complexes on the beads were incubated with 40 units T4 Polynucleotide kinase (New England Biolabs), first with P32 labelled ATP for 45 min, then 20 more min with 1 mM cold ATP, in PNK buffer with RNase inhibitors (RNasin, Promega) at room temperature. The reaction should provide 5’ phosphates needed for downstream ligations. The beads then were washed as before twice UB (FLASH) or GDB (CLASH) and 4x PNK buffer.

### Linker ligation and RNA-protein complex elution

Protein-bound RNA molecules were ligated together and with 3’ linker (1 μM miRCat-33, IDT), overnight using 40 units of T4 RNA ligase 1 (New England Biolabs) in PNK buffer with RNase inhibitors at 16°C. This reaction created RNA hybrids and single RNA molecules ligated to miRCat linker. On the next day, the beads were washed as before 2x UB (FLASH) or GDB (CLASH) and 4x PNK buffer. Then using 40 units of RNA ligase 1, barcoded 5′ linkers (final conc. 5 μM; IDT, one for each sample) were ligated in RNA ligase 1 buffer with 1mM ATP for 3-6 h at 20°C. The beads were washed as before. In CLASH the complexes were eluted in EB (2x NuPage Sample buffer, 400 mM Imidazole, 10mM Tris-HCl pH=7.4, 10mM DTT). In FLASH the complexes were washed off the beads by partial destruction of formaldehyde crosslinking by boiling the samples in NuPAGE protein sample buffer plus 100 mM Tris-HCl, 1%SDS, 100 mM ME (β-mercaptoethanol) for 3 min. The supernatant with RNA-protein complexes was recovered from cooled samples.

### SDS-PAGE, and transfer to nitrocellulose

Protein-RNA complexes in NuPAGE SB plus SDS, ME (Life Technologies) were resolved on a 4%–12% Bis-Tris NuPAGE gel (Life Technologies) in NuPAGE SDS MOPS running buffer then they were transferred to nitrocellulose membrane (GE Healthcare, Amersham Hybond ECL) in NuPage transfer buffer (Life Technologies) with 10% methanol for 1 hr at 100V. Depending on the strength of the signal the membrane was exposed on film (Amersham) for 1 hr or overnight at -70C. Developed film was aligned with the membrane and the radioactive bands corresponding to the Fibrillarin-RNA complexes were excised.

#### Proteinase K Treatment, RNA Isolation and cDNA Library preparation

Cut out bands were incubated with 150 μg of Proteinase K (Roche) and proteinase K buffer (50 mM Tris-HCl pH 7.8, 50 mM NaCl, 0.4% NP-40, 0.5% SDS, 5 mM EDTA) for 2 hr at 55°C. The

RNA was extracted with phenol-chloroform-isoamyl alcohol (PCI) mixture and ethanol precipitated overnight with 10 μg Glycogen (Ambion, Life Technologies). The isolated RNA was dissolved in 12 mkL of distilled RNAse-free water and reverse transcribed using miRCat-33 primer (IDT) with Superscript III Reverse Transcriptase (Life Technologies) in its buffer for 1h at 50°C. RNA was then degraded by addition of RNase H (New England Biolabs) for 30 min at 37°C. cDNA was amplified using primers P5 and primer PE_miRCat_PCR and TaKaRa LA Taq polymerase (Takara Bio). PCR products were separated on a 2% MetaPhor agarose (Lonza) gel with SYBRSafe (Life Technologies) in 1 x TBE at +4C. The gel band corresponding to 150- 200bp was cut out. cDNA was purified with MinElute Gel Extraction Kit (QIAGEN). Obtained cDNA libraries were sent for high-throughput sequencing.

### Oligonucleotides 3’ linker

miRCat-33 linker (IDT) AppTGGAATTCTCGGGTGCCAAG/ddC/

### 5’ linkers

L5Aa invddT-ACACrGrArCrGrCrUrCrUrUrCrCrGrArUrCrUrNrNrNr<u>UrArArGrC</u>-OH

L5Ab invddT- ACACrGrArCrGrCrUrCrUrUrCrCrGrArUrCrUrNrNrN<u>rArUrUrArGrC</u>-OH

L5Ac invddT-ACACrGrArCrGrCrUrCrUrUrCrCrGrArUrCrUrNrNrN<u>rGrCrGrCrArGrC</u>-OH

L5Bb invddT-ACACrGrArCrGrCrUrCrUrUrCrCrGrArUrCrUrNrNrN<u>rGrUrGrArGrC</u>-OH

L5Bc invddT-ACACrGrArCrGrCrUrCrUrUrCrCrGrArUrCrUrNrNrN<u>rCrArCrUrArGrC</u>-OH

L5Bd invddT-ACACrGrArCrGrCrUrCrUrUrCrCrGrArUrCrUrNrNrN<u>rUrCrUrCrUrArGrC</u>-OH

L5Ca invddT-ACACrGrArCrGrCrUrCrUrUrCrCrGrArUrCrUrNrNrN<u>rCrUrArGrC</u>-OH

L5Cb invddT-ACACrGrArCrGrCrUrCrUrUrCrCrGrArUrCrUrNrNrN<u>rGrGrArGrC</u>-OH

L5Cc invddT-ACACrGrArCrGrCrUrCrUrUrCrCrGrArUrCrUrNrNrN<u>rArCrTrCrArGrC</u>-OH

L5Cd invddT-ACACrGrArCrGrCrUrCrUrUrCrCrGrArUrCrUrNrNrN<u>rGrArCrTrTrArGrC</u>-OH

The fixed barcodes are underlined. N indicates mixed nucleotides for random barcodes.

### PCR primers

miRCat-33 primer (IDT) CCTTGGCACCCGAGAATT primer for RT

PE_miRCat_PCR

CAAGCAGAAGACGGCATACGAGATCGGTCTCGGCATTCCTGGCCTTGGCACCCGAGAATT

CC library amplification

P5 AATGATACGGCGACCACCGAGATCTACACTCTTTCCCTACACGACGCTCTTCCGATCT

library amplification

### Bioinformatics

#### Analysis of CLASH data

Raw sequences were preprocessed prior to alignment using hyb (Travis et al., 2014) by running the hyb preprocess command with standard parameters. The preprocessed data were aligned to a custom database combining multi-exon transcripts and unspliced genes (with snoRNA genes extended by 20bps in each direction and masked out of the genes in which they are contained where appropriate). The custom database was built using reference data from Ensembl release 77 (www.ensembl.org). To facilitate the analysis of snoRNA/rRNA hybrids, the complete human ribosomal DNA repeating unit (https://www.ncbi.nlm.nih.gov/nuccore/U13369) was also included in the database. Sequence alignment was performed using the blastall command, using the standard parameters from the hyb pipeline (Travis et al., 2014). The aligned reads were processed using a variant of the hyb pipeline, modified slightly to extract snoRNA hybrids rather than microRNA hybrids preferentially. Hybrids identified using this process were then filtered to exclude sequences that could be aligned as single reads to the human genome (Ensembl release 77) using Novoalign 2.07 (www.novocraft.com) to prevent single reads overlapping gene boundaries from being mistakenly identified as hybrids. Downstream analysis was performed on reproducible hybrids (in which both fragments were found to overlap in two or more hybrids) with a predicted folding energy of -12dG or below. Among these stable, reproducible hybrids, further filters were applied to ensure that hybrids between snoRNAs and other classes of RNA were not mismapped U3 stems, and that hybrids between snoRNAs and RNAs that were not snoRNAs or rRNAs were not mismapped snoRNA-rRNA hybrids. The analysis was performed using the hybtools python package (http://www.github.com/hyweldd/hybtools), which was developed for this project. Reference data for the analysis of human methylation sites were obtained from (Krogh et al., 2016) and (Jorjani et al., 2016).

All interactions recovered are listed in Dataset 1.

#### Analysis of RNA-Seq data

RNA-Seq data were processed using STAR (Dobin et al., 2013) and DESeq2 (Love et al., 2014), using a human genome database from ensemble release 77 (www.ensembl.org).

## RESULTS

### Systematic mapping of box C/D snoRNA interactions by UV crosslinking

To identify potential novel snoRNA interactions we initially applied the CLASH technique (Figure 1A) (Helwak et al., 2013; Kudla et al., 2011). This involves UV crosslinking of RNA complexes with tagged proteins in living cells and affinity purification of the RNP complexes under stringent conditions. Ligation of linker adaptors is performed in parallel with internal ligation of captured RNA fragments base paired to each other. RNA is isolated, including RNA hybrids, followed by reverse transcription and high throughput sequencing of cDNA libraries.

**Figure 1.**
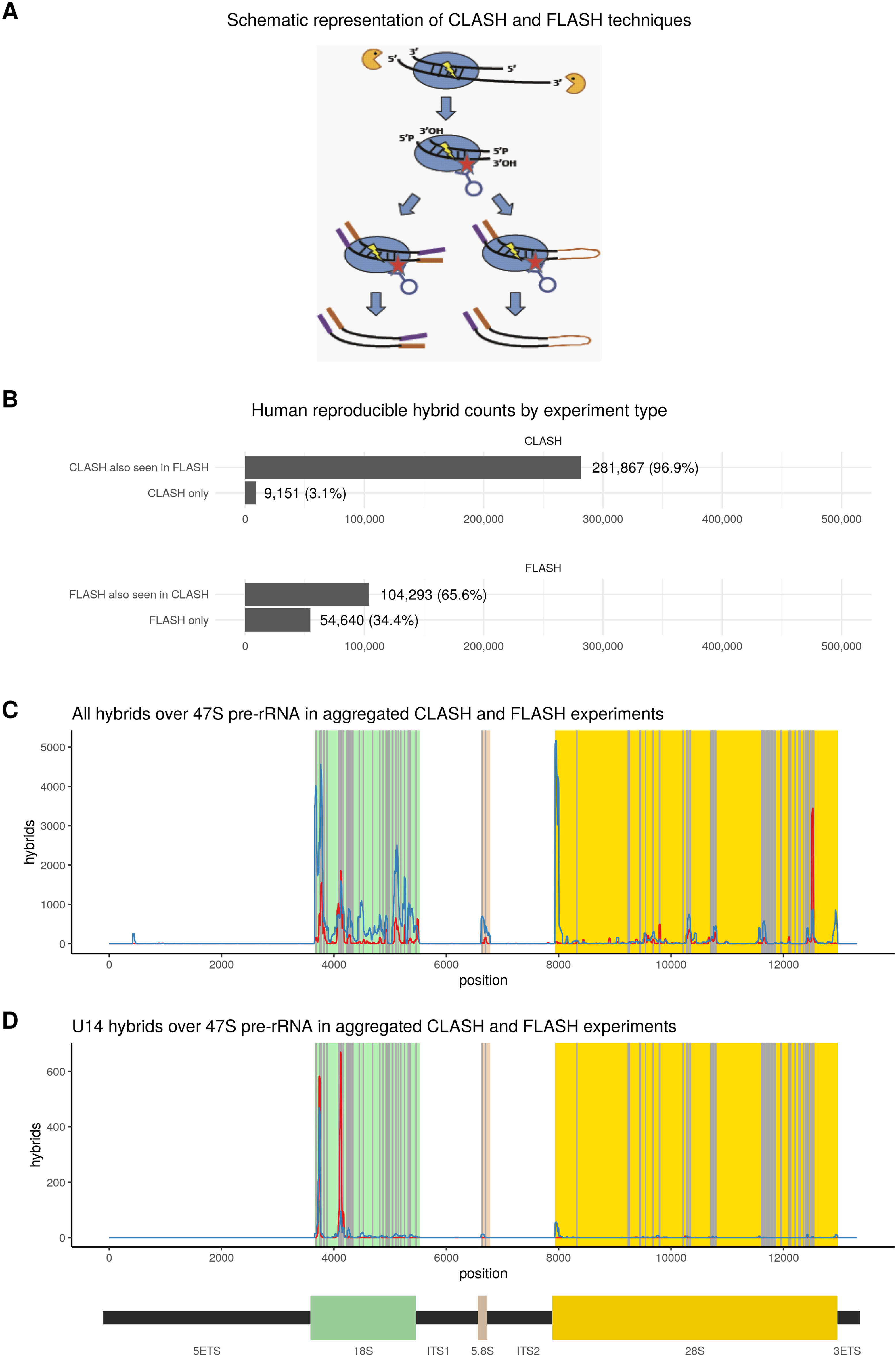
Overview of CLASH and FLASH results. (A) Schematic representation of CLASH and FLASH techniques. Live cells were UV-irradiated, crosslinked RNA–protein complexes were extracted and affinity purified. RNAs were partially digested and interacting RNAs were ligated together to generate hybrid molecules. Following cDNA generation and sequencing, cognate RNA pairs were identified in chimeric cDNAs. In CLASH, complexes were tandem-affinity purified under denaturing conditions. In FLASH, complexes were immuno-purified using antibodies and then crosslinked to the beads, allowing subsequent wash steps under denaturing conditions. (B) Human reproducible hybrid counts by experiment type. Reproducibility of hybrids across CLASH and FLASH experiments in HEK293 cells. (C) All hybrids over 47S pre-rRNA in aggregated CLASH and FLASH experiments. The distributions of snoRNA hybrid hits in CLASH (red) and FLASH (blue) over 47S pre-rRNA show similar peaks, with known methylation sites shown in grey. (D) U14 hybrids over 47S pre-rRNA in aggregated CLASH and FLASH experiments. U14 (SNORD14) hybrids with rRNA in CLASH (red) and FLASH (blue) identify known interaction sites.

CLASH analyses require the use of a “bait” protein fused to a tandem affinity purification tag. For these analyses, we tagged the core box C/D snoRNP proteins Fibrillarin (FBL) or NOP56 with a C-terminal tag consisting of His6 – Precision protease cleavage site – Flag epitope. Tagged constructs were integrated into the chromosome in Flip-in HEK293 cells at a pre-inserted LoxP site. The fusion proteins were expressed under a regulated pCMV-2xTET O2 promoter.

Tetracyclin levels were titrated to achieve expression close to the endogenous protein level, as assessed by western blotting. The tagged proteins were previously shown to be functional in rRNA processing by rescue experiments, in which the endogenous proteins were depleted by RNAi (Knox et al., 2011).

In order to assess robustness of our dataset we performed orthogonal validation, using formaldehyde assisted crosslinking ligation <u>a</u>nd <u>s</u>equencing of <u>h</u>ybrids (FLASH) (Bharathavikru et al., 2017). This approach is similar to CLASH in that RNA-protein interactions are captured by UV crosslinking in growing cells, but antibodies are used for affinity purification of endogenous RNA-protein complexes. During purification, brief formaldehyde crosslinking is used to stabilize binding of the covalent bait protein-RNA complex to the protein A beads, allowing column washes under highly denaturing conditions. Analyses of single hits for FBL and NOP56 showed that, for both proteins, snoRNA sites were most frequently recovered followed by rRNA and then mRNA hits, in both CLASH and FLASH analyses (Supplementary Figure 1), supporting the reliability of the crosslinking approaches.

In human cells we recovered 591,958 hybrids overall (Supplementary Figure 2A; Table S1; Dataset 1). Recovered sequences that could be confidently mapped to two distinct regions of the genome (see Methods) were regarded as representing chimeric cDNAs resulting from RNA-RNA ligation. Non-identical chimeric sequences, or sequences recovered from different analyses, in which both segments overlapped were regarded as demonstrating independent recovery of the same interaction. The recovered RNA sequences were folded *in silico,* using the ViennaRNA Package 2.0 (Lorenz et al., 2011), to assess whether they arose from a stable RNA-RNA duplex. Interactions supported by at least two independent sequences, with a predicted ΔG of less than -12 Kcal mol^-1^, were considered stable and reproducible, and included in further analyses; this was the case for a total of 449,781 hybrids (Supplementary Figure 2A). Among stable, reproducible hybrids, further filters were applied to ensure that hybrids called between snoRNAs and other classes of RNA were not mismapped internal snoRNA stems, and that hybrids between snoRNAs and RNAs that were not snoRNAs or rRNAs were not mismapped snoRNA-rRNA hybrids.

We compared CLASH from the cells expressing tagged Fibrillarin with FLASH from untagged control cells using anti-Fibrillarin antibodies. Strikingly, 97% of stable, reproducible RNA-RNA interactions recovered by CLASH were mapped to sites of interactions also recovered in FLASH: (Figure 1B). A lower fraction of hybrids recovered by FLASH corresponded to interaction sites also found in CLASH data (66% of hybrids), with 34% FLASH only hybrids (Figure 1B).

The majority of hybrids mapping to snoRNAs were internal, representing stem structures (Supplementary Figure 2; Table S1). These potentially allow visualization and analysis of snoRNA structures. Among intermolecular snoRNA hybrids, snoRNA-rRNA hybrids were most frequently recovered. From the set of stable, reproducible intermolecular hybrids after filtering, 69% were snoRNA-rRNA interactions, 9% were snoRNA-mRNA interactions, and 17% were snoRNA-snoRNA. It is notable that some highly abundant RNA species were recovered at low levels, in particular snoRNA-tRNA interactions represented only 0.7% of intermolecular hybrids, supporting the specificity of the interactions. The predominant recovery of snoRNA-rRNA interactions is consistent with the known function of snoRNAs in ribosome synthesis. Recovery of different RNA species in single reads (Supplementary Figure 1) and chimeras (Supplementary Figure 2) was in general agreement, supporting the recovery of authentic interactions.

To confirm the reliability of both methods we compared snoRNA-rRNA interactions recovered as hybrids in both types of experiments with the position of known rRNA methylation sites (Figure 1C). Comparing CLASH and FLASH results for snoRNA-rRNA targeting in human HEK cells we noticed that although peak intensities varied to some extent, the same major interactions were recovered with both methods and correlated with known rRNA methylation sites. This was strongly supported by analyses of individual snoRNA interactions e.g. U14, which is known to interact at two positions on the 18S rRNA sequence (Figure 1D). We conclude that both CLASH and FLASH provide consistent and reliable results. However, the background in FLASH analyses appeared higher than with CLASH, presumably reflecting the lower stringency of purification.

### Identification of novel snoRNA-rRNA interaction sites

Modification sites in ribosomal RNAs have been well characterized by a variety of highly sensitive techniques, and it is very likely that all high-efficiency methylation sites have been identified (see (Marchand et al., 2016; Taoka et al., 2016) and references therein). Nonetheless, sub-stoichiometric modifications may have escaped direct identification. If these are mediated by a box C/D snoRNA, a hybrid must form that may be captured by CLASH. We therefore developed robust bioinformatics filters for snoRNA-rRNA interactions, to confidently identify putative novel sites of methylation.

We use the following strict filtering criteria to identify interactions that we can classify with high confidence as being capable of guiding methylation (Figure 2):

- Nucleotide 5 bps upstream of D box or D’ box in snoRNA base pairs exactly with rRNA and hybrid contains 12 or more base-paired nucleotides
- One or more nucleotides in region between 2 and 4 bps upstream of D box or D’ box base pairs exactly with rRNA
- There is not exact base pairing over the whole D or D’ box
- Either:

- There is a stretch of eight nucleotides between 1 and 14 bps upstream of D box or D’ box base pair exactly with rRNA (no mismatches), or
- There is a stretch of eleven nucleotides between 1 and 16 bps upstream of D box or D’ box base pair exactly with rRNA with at most 1 mismatch

**Figure 2.**
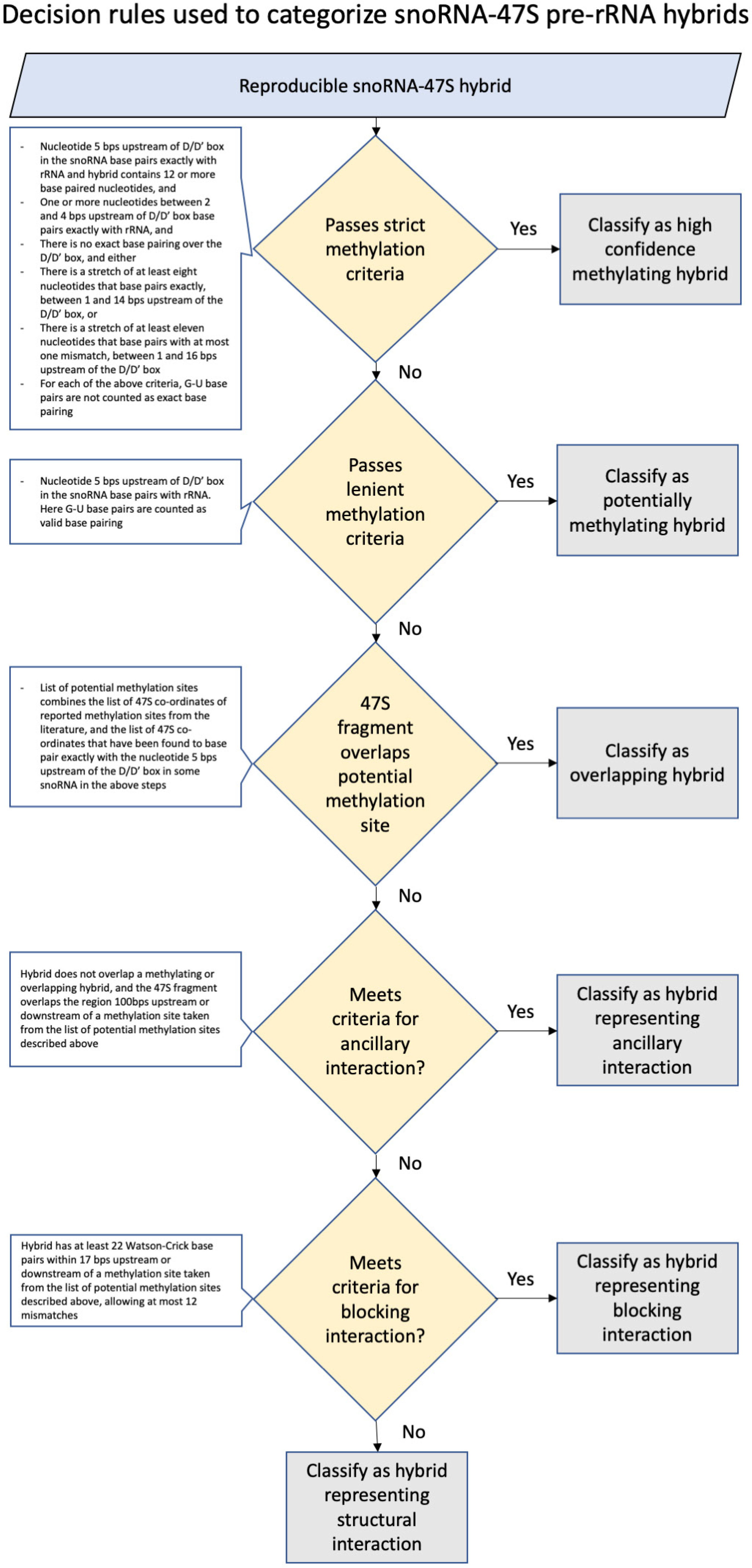
Decision rules used to categorize snoRNA-47S pre-rRNA hybrids. The flowchart summarizes the decision rules. The filtering criteria were applied to reproducible snoRNA-rRNA hybrids to identify those that are likely to guide methylation.

For each of the above criteria, G-U base pairs are not counted as exact base pairing.

We recovered 9,363 hybrids passing these criteria and thus potentially able to guide methylation In addition to known methylation sites we recovered novel snoRNA-rRNA interactions that could potentially guide methylation, including an orphan snoRNA SNORD126 (Figure 3; Supplementary Table S2). For SNORD14 (U14) interactions were recovered that would potentially guide methylation at 18S-462 and 18S-83. We conclude that the filters are quite conservative, and should recover only interactions with a high likelihood of representing methylation-guide RNA binding sites.

**Figure 3.**
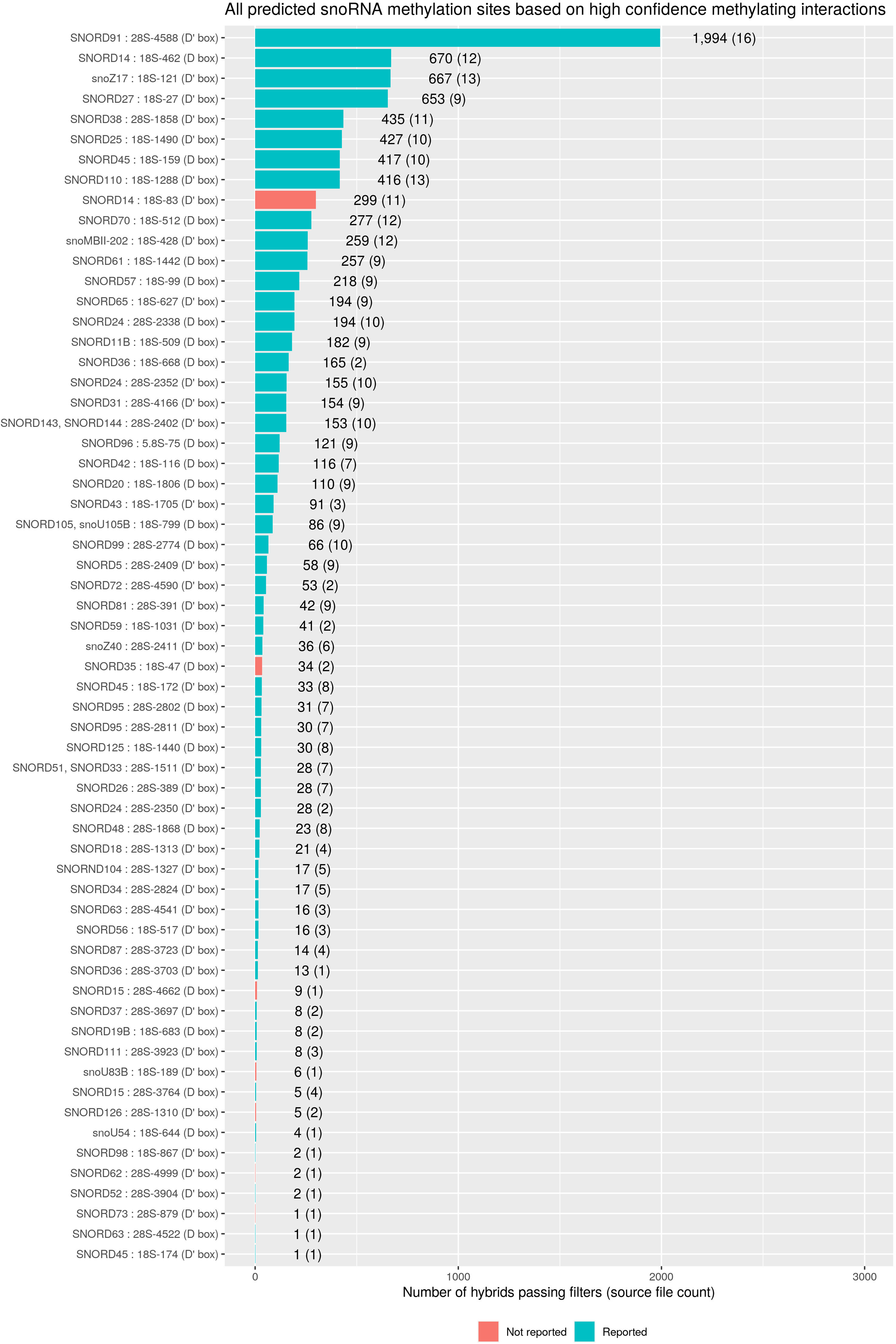
Distribution of predicted cognate snoRNA-rRNA interactions at methylation sites. Predicted sites of C/D box snoRNA guided methylation based on the filtered set of reproducible snoRNA-rRNA hybrids. The x axis labels show the snoRNA, the pre47S subunit and co-ordinate at which methylation is predicted to occur, and the snoRNA box 5 base pairs downstream of the methylating nucleotide. The bars show the number of hybrids associated with the methylation event described in the access label, with the number of experiments in which relevant hybrids were found shown in brackets. Blue bars indicate previously reported methylation sites, and red bars indicate interactions that have not previously been reported to be associated with methylation.

Due to the strict criteria, many confirmed methylation site interactions were filtered out. In order to ensure that hybrids that did not meet these strict criteria but were associated with methylation site interactions were not included in downstream analyses, we also used a weaker set of criteria to identify potentially methylating hybrids. Hybrids were classified as potentially methylating hybrids if they did not meet the strict methylation criteria, but the nucleotide 5 base pairs upstream of the D or D’ box did base pair with the rRNA. Unlike the strict criteria, these weaker criteria did allow G-U base pairs. We recovered 7,971 hybrids passing these criteria, representing 412 interactions, including additional known methylation guide interactions for which no hybrids were found that passed the strict methylation criteria (Table S3).

The hybrids that did not meet the strict or lenient methylation criteria, and whose pre-47S fragment did not overlap with a nucleotide for which the snoRNA included in the hybrid was known to guide methylation, were separated in three further categories: ‘ancillary’ (Table S4), ‘blocking’ (Table S5), or ‘structural’ hybrids (Table S6), depending on whether they potentially assist or interfere with the methylation function of a snoRNA, or contribute to pre-rRNA folding during ribosomal subunit assembly (Figure 2).

Hybrids recovered 100 nt upstream or downstream of a methylation site directed by the same snoRNA, but not overlapping with it, were designated “ancillary” as they could give additional structural support to the guide snoRNA interaction (Figure S3A; Supplementary Table S1) (van Nues et al., 2011). We recovered 692 ancillary hybrids.

Hybrids formed by snoRNAs at methylation site, that are not predicted to guide methylation but forming at least 22 perfect Watson-Crick pairs within 17 nt upstream or downstream of the site were called “blocking” interactions. In total, 11,028 non-methylating hybrids overlapped with interactions guiding methylation (Supplementary Table S1). Among these interactions, the majority guide methylation at neighboring sites. It is unlikely that closely located sites (less than 20 nt separation) could be methylated. Thus, over-expression of a snoRNA could lead to both increased methylation at its target rRNA site and suppression at neighboring sites. The presence of such overlapping methylation guide interaction sites suggests the need for a precise timing for snoRNA binding and methylation; such an ordered sequence may contribute to the correct folding of the pre-rRNAs and/or aid in avoiding kinetic traps (Huang and Karbstein, 2021; Steitz and Tycowski, 1995). For clarity, these interactions were not included in the “blocking” interactions list. However, 62 high confidence blocking hybrids were identified for snoRNAs that are not predicted to direct methylation at closely located sites (Figure S3B; Supplementary Table S1). Recent reports have highlighted the variability in methylation efficiency at different sites in the human rRNA (Erales et al., 2017; Krogh et al., 2016; Sharma et al., 2017a; Zhou et al., 2017):, and we speculate that this may partly reflect competition for binding between snoRNA species. For instance, we observed interactions involving the abundant snoRNAs U3 and U8 that could block methylation sites.

All other confidently identified snoRNA-rRNA hybrids were termed “structural” interactions, reflecting potential structural roles in supporting conformational changes and avoiding kinetic traps during pre-ribosome assembly and/or pre-rRNA folding (Huang and Karbstein, 2021; Steitz and Tycowski, 1995). These included the small number of box C/D snoRNAs implicated in ribosome synthesis steps other than rRNA methylation (Figure S3C): U3, U8, U14 and the acetylation guide U13.

Among novel structural interactions, we found hybrids between 18S rRNA and the 3’ region of U8 snoRNA. It was previously reported that the 5’ end of U8 snoRNA is critical for 5.8S and 28S rRNA maturation (Peculis, 1997). However, no interactions for the 3’ region of U8 were described and the functional significance remains to be established. In *Xenopus*, the timing of association of the 3’ end of 5.8S rRNA and 5’ end of 28S was proposed to be regulated by initial binding of U8 at the 5’ end of 28S, promote formation of a “bulge” in the 28S sequence. This might act as a “priming site” for base-pairing to 5.8S, leading to the eventual displacement of U8. We note that a peak of U8 interaction is located at the 5’ end of 28S (Figure 4B), potentially corresponding to this predicted interaction.

**Figure 4.**
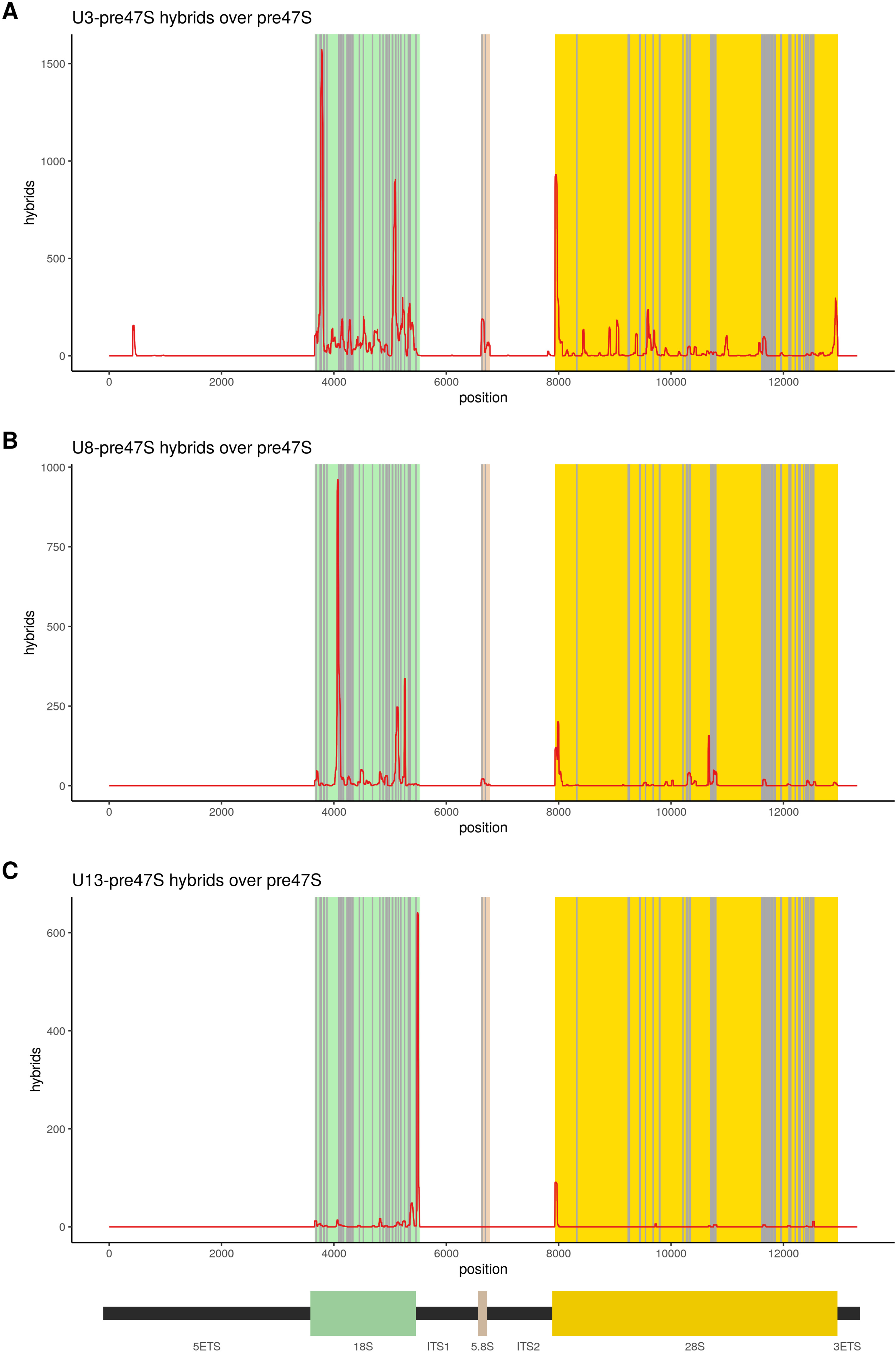
Distribution of selected snoRNAs on the 47S pre-rRNA. (A-C) The distribution of hybrid hits involving U3, U8, and U13, respectively, over the 47S pre- rRNA sequence. Reported methylation sites are shown in grey.

### Novel snoRNA-snoRNA interactions

Many reproducible snoRNA-snoRNA hybrids were recovered (261,166) (Supplementary Table S1), but these predominately represented internal stems, particularly within U3 (229,525 hybrids). However, we also recovered 14,235 reproducible intermolecular hybrids between different snoRNA species, representing 1,422 distinct interactions (Figures S4A and S4B).

These interactions suggest the existence of regulatory loops in snoRNA biogenesis and function, e.g. through possible titration/sequestration or “sponging”. In addition, a small number of interactions predicted to guide snoRNA methylation were detected (Figure S4C), using the same criteria as applied to rRNA (Figure 2).

### snoRNA-tRNA interactions

We noted that although tRNAs represented only a small proportion of all snoRNA hybrids (0.2%), they were enriched for interactions that potentially direct methylation. Overall, from 591 reproducible snoRNA-tRNA hybrids, two met the criteria for classification as high confidence methylating hybrids, and 161 met the criteria for classification as potentially methylating hybrids. Notably, for hybrids between snoRNAs and tRNAs that contain introns, 63% (71 out of 113 reproducible hybrids) were classified as high confidence or potentially directing methylation. In contrast, for hybrids between snoRNAs and tRNAs that do not contain introns, only 19% (92 of 478 reproducible hybrids) meet these criteria (Supplementary Table 1).

### snoRNA-mRNA interactions

Perhaps the most interesting class of snoRNA chimeras involved snoRNA-mRNA interactions. It has been proposed that snoRNAs can influence pre-mRNA splicing, processing and stability in mammalian cells; for examples see (Falaleeva et al., 2015; Huang et al., 2017; Kishore and Stamm, 2006; Sharma et al., 2016). It is, however, also possible that snoRNAs might be sponged on abundant mRNAs.

Among reproducible, stable hybrids, 28,120 snoRNA-mRNA hybrids were recovered, representing 1,755 interactions between 149 snoRNA families and 967 mRNAs. To eliminate potential mis-mapping errors we removed all hybrids that were called as snoRNA-mRNA hybrids, but whose mRNA fragment could also be aligned to U3 or to an rRNA sequence (albeit poorly). This filtering step retained 7,209 hybrids involving 117 snoRNAs and 940 mRNAs.

The greatest number of filtered mRNA interactions was observed for U3 (50% of reproducible snoRNA-mRNA hybrids, interacting with 566 different mRNAs), followed by SNORD33 (8% of hybrids, 23 mRNAs), SNORD24 (3% of hybrids, 15 mRNAs), snoU83B (3% of hybrids, 63 mRNAs), and SNORD58 (3% of hybrids, 37 mRNAs). There was a clear correlation for mRNAs between filtered hybrids and single hits (p = 6e^−166^), showing enrichment of mRNA single hits in the regions of snoRNA interactions (Figure 5B). In contrast, there was little correlation between mRNA expression levels and recovery in snoRNA hybrids (R^2^=0.0057) (Figure 5C). These data support the conclusion that target mRNAs were specifically recovered and represent *bona fide* interactions.

**Figure 5.**
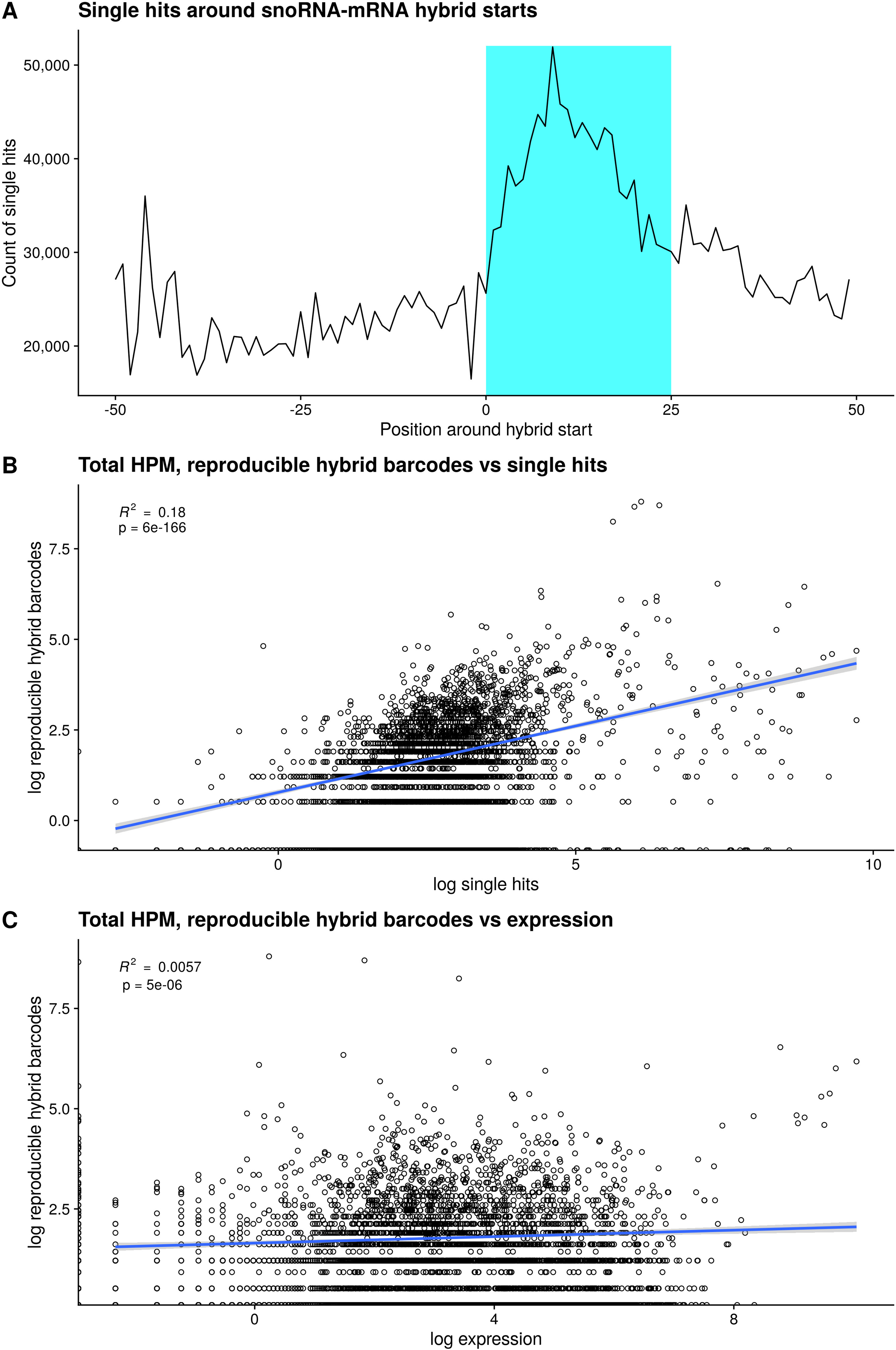
Single hits around snoRNA-mRNA hybrid starts, and correlation of snoRNA-mRNA hybrids with mRNA single hits and abundance (A) Distribution of single hits around snoRNA-mRNA hybrid starts. (B) Correlation between recovery in reproducible CLASH hybrids and mRNA single hits. (C) Correlation between recovery in reproducible CLASH hybrids and mRNA abundance.

To assess whether novel snoRNA-mRNA interactions have the potential to direct RNA methylation, the hybrids were analyzed using the same criteria those applied to rRNA (Figure 2). This identified a small number of putative methylation guide interactions (Supplementary Figure S7).

Comparison of snoRNA-mRNA interactions revealed distinctly different patterns of targets between U3 and other snoRNAs (Figure 6A). Interactions with all snoRNAs were recovered in mRNA coding sequences (CDS) and untranslated regions (UTRs), but were substantially enriched in pre-mRNA introns (Figure 6A). However, more CDS interactions were recovered for U3 than for all other snoRNAs combined. Those U3 interactions that were identified within mRNA introns were predominately not in proximity to splice junctions. Moreover, most mRNA-U3 binding sites presented sharp peaks pointing to highly-specific interactions. The strikingly high number of U3-mRNA interactions suggest a special role for U3 in mammalian gene expression, which might be reflected in the substantially greater abundance of U3 than other human snoRNAs and detection of stable abundant U3-derived fragments.

**Figure 6.**
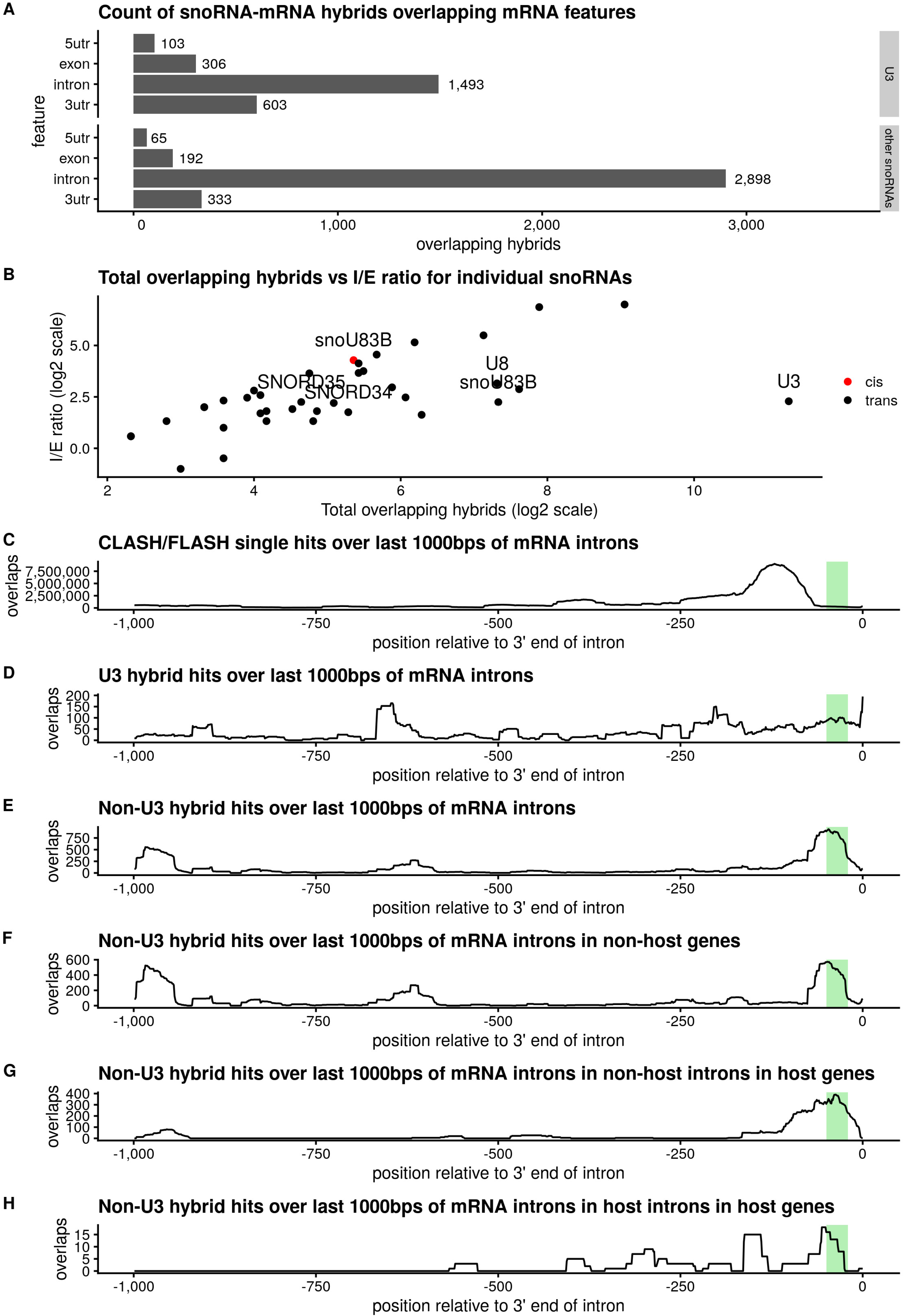
Distribution of snoRNA interactions on mRNAs. (A) Counts of hybrids overlapping different mRNA features for snoRNA−mRNA hybrids incorporating U3, and hybrids incorporating other snoRNAs, respectively. The plots show that while both classes of hybrid predominantly bind mRNA introns, hybrids incorporating U3 are significantly more likely to bind other regions of their target RNA. (B) Total overlapping hybrids vs intron / exon ratios for individual snoRNAs in cis and trans. (C-H) CLASH / FLASH hits over the last 1000 base pairs of mRNA introns, with the estimated branch point region (20-50bps upstream of the 3’ end of the intron) highlighted in green. Non-U3 hybrids show a pronounced peak overlapping the estimated branch point region in host introns, non-host introns within the host gene, and introns within non-host genes.

To study the functional role of U3 interactions we depleted U3 in HEK cells with morpholino oligo and carried out transcriptome analyses of control (mock treated) and U3-depleted cells. RNA sequencing showed that 3 days following U3 depletion, substantially more mRNAs identified as U3 targets in CLASH showed altered levels than non-targets or total genes (Chi Square Test; p = 4e^-6^). U3 target RNAs showed both increased and decreased levels, but 20% of CLASH/FLASH U3 targets showed reduced abundance after U3 depletion (Supplementary Figure S8B). Notably, we observed a correlation between the presence of a snoRNA targeted by U3 in the intron and levels of its host mRNA after U3 depletion, pointing to general rule of interplay between biogenesis of snoRNAs and their host genes.

Initial comparison of the binding sites to the patterns of evolutionary conservation across 23 mammalian species did not identify enrichment for conserved regions in snoRNA binding sites relative to their flanking regions. However, further analysis revealed that the fall in conservation was due to the frequent presence of an exon near the snoRNA binding site. The 3’ region of introns that bind snoRNAs was more conserved than, for instance, the 5’ region of the same intron not harboring interactions. This observation supports the functional importance of the interactions (Supplementary Figure S6).

### Binding is frequently found within introns that host snoRNAs

We noted that a substantial proportion of snoRNA hybrids with intronic regions represented interactions between snoRNAs and their host introns (5% or 220 hybrids), suggesting frequent connections between snoRNA biogenesis and host gene splicing (Figure 6). Around 16% of all C/D snoRNA-mRNA interactions were represented by peaks towards the 3’ ends of introns, *in cis* or *in trans,* in the region of potential intron branch point (25 to 50 bp from the 3’ SS) (Figure 6) pointing to the possibility of their involvement in splicing of the target mRNAs.

Intronic snoRNAs frequently formed predicted duplexes with the 3’ flanking region that included the intron branch point and/or of the polypyrimidine tract, both of which are important signals for pre-mRNA splicing. For examples see Figure 7. We speculate that these interactions may slow pre-mRNA splicing allowing sufficient time for assembly of snoRNP proteins with the nascent transcript prior to pre-mRNA intron cleavage and degradation.

**Figure 7.**
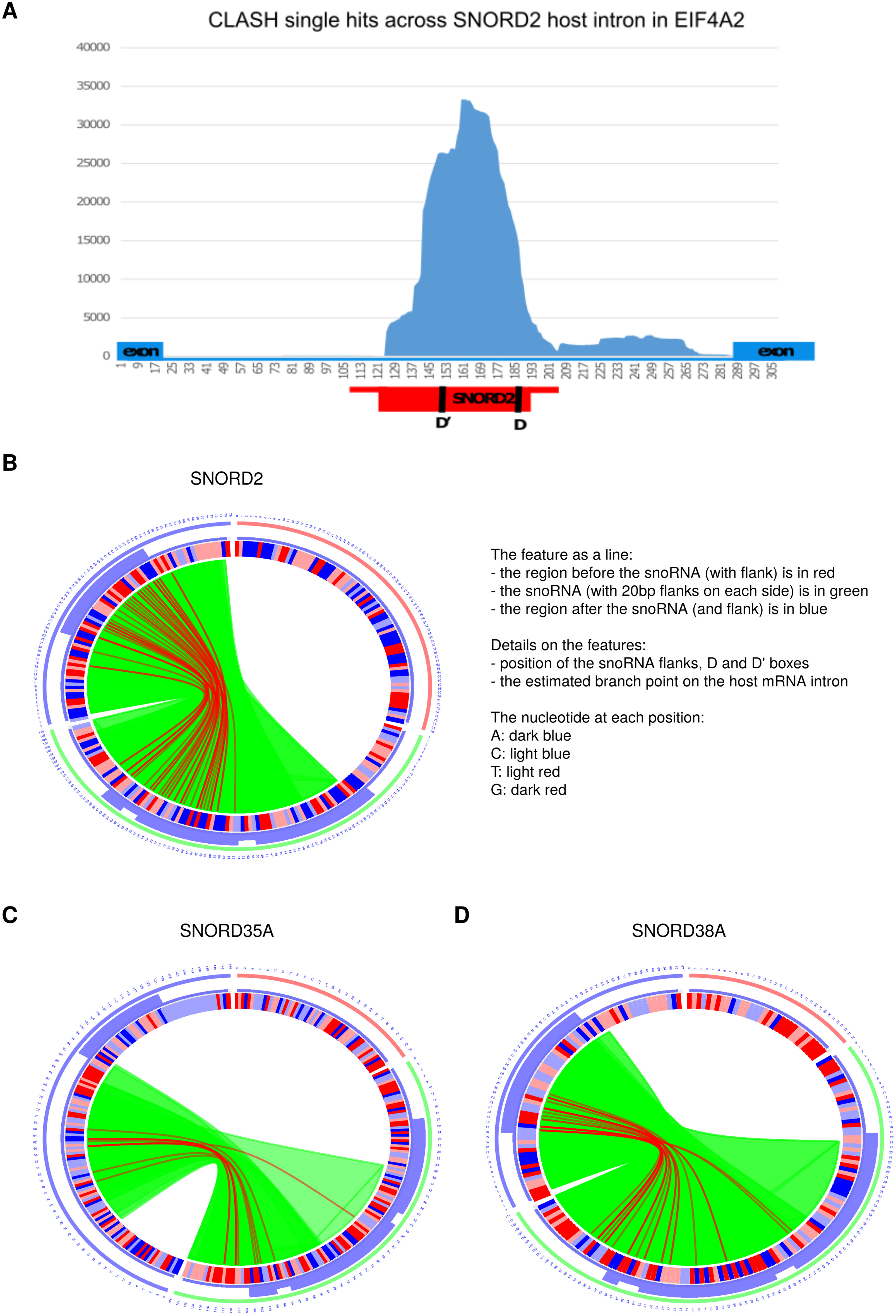
Interactions between selected snoRNAs and host introns. (A) Distribution of CLASH single hits across the host intron of SNORD2 in EIF4A2, showing that reads that do not overlap the snoRNA overlap the 3’ end of its host intron (B,C) Circos plots showing hybrids between SNORD2, SNORD35A, and SNORD38A, and their host introns. In each case, hybrids are exclusively formed between the snoRNA and the 3’ end of its host intron.

## DISCUSSION

Here we systematically mapped the interactions between box C/D snoRNAs and the human transcriptome by the use of UV crosslinking, followed by generation and sequencing of RNA hybrids. To achieve high specificity and robustness, we used two complementary approaches: CLASH relies on high-affinity purification tags, suitable for extensive washes in denaturing conditions, whereas FLASH utilizes specific antibodies in combination with mild chemical crosslinking with formaldehyde to stabilize the interactions. Comparison of snoRNA hybrids recovered with CLASH and FLASH revealed a high degree of overlap, and we therefore combined these datasets for most analyses.

As expected, we recovered many interactions between the cognate snoRNAs and known or predicted sites of rRNA methylation. It was, however, notable that at many sites of methylation the human rRNA, multiple snoRNAs could be identified with complementary base-pairing that matched the features previously defined as required for 2’-*O*-methylation; ≥12 bp showing perfect complementarity, with the modified nucleotide positioned 5 bp from a site D motif (Kiss-László et al., 1996). In several cases previously predicted snoRNAs were not recovered, although these may well bind under altered metabolic conditions, in different cells types or during cell differentiation and tumorigenesis. These data underline the surprisingly high degree of plasticity and redundancy in human snoRNA-rRNA interactions. Recent quantitative analyses of human rRNA methylation have identified many sites that show partial methylation suggesting that functionally distinct ribosomes are generated under different growth conditions (Erales et al., 2017; Hebras et al., 2020; Krogh et al., 2016; Sharma et al., 2017a). We also found numerous cases in which snoRNAs were recovered bound to the pre-rRNA close to methylation sites with strong Watson-Crick base-pairing, but in a configuration that is not expected to guide RNA modification. We speculate that these interactions regulate the timing and/or efficiency of rRNA modification by competing with cognate methylation-guide snoRNAs. In addition, they may contribute to pre-rRNA folding dynamics during ribosomal subunit biogenesis, as previously proposed (Huang and Karbstein, 2021; Steitz and Tycowski, 1995). The emergence of snoRNAs with overlapping specificities and overlapping binding sites, indicate that the site-specific regulation of rRNA methylation is both functionally important and complex.

A large number of reproducible snoRNA-mRNA hybrids represented snoRNA-mRNA interactions. Human U3 is 20-50 fold more abundant than most methylation-guide snoRNAs and was responsible for 50% of all filtered snoRNA-mRNA hybrids. These interactions involved a 3’ sequence in U3, homologous to a guide region in yeast U3 previously implicated in non-canonical interactions (Dudnakova et al., 2018; Kudla et al., 2011). Following U3 depletion, mRNAs identified as bound by U3 were highly over-represented among mRNAs showing altered abundance. Moreover, U3 target sites were largely located in exons, whereas other snoRNAs predominately target conserved regions of pre-mRNA introns. However, the mechanistic links between U3 interactions and altered mRNA abundance remain unclear. Notably, snoRNA-mRNA interactions did not resemble those formed by miRNAs in structure, distribution or, most likely, in function. The length distribution of the snoRNA fragment of snoRNA-mRNA chimeras showed an average length of around 35 to 40-nt with base pairing region of at least 12nt. This is much longer than expected for miRNA interactions. We also did not observe accumulation of the previously reported, short (≤ 22nt), miRNA-like fragments of snoRNAs in single reads.

In human cells, the majority of snoRNA genes are located within introns in protein coding genes. It was previously suggested that processing of snoRNAs and splicing of the host gene may be connected (Hirose et al., 2003). Indeed, folding of the snoRNA snoRD86, which is encoded in an intron of the NOP56 gene, acts as sensor in controlling the abundance of this snoRNP core protein (Lykke-Andersen et al., 2018). We observed that intronic regions flanking snoRNAs were frequently recovered in hybrids with the same snoRNA, as well as with other snoRNAs. We suggest that such interactions facilitate coordination between the maturation of snoRNA and splicing of the host gene. Debranched introns are expected to be rapidly degraded by both the 5’ exonuclease Xrn2 and 3’ exonucleases of the exosome complex. It is therefore important that snoRNA folding and snoRNP assembly precedes intron excision. The observed interactions may coordinate these steps.

Despite the essential roles they play during ribosome biogenesis through their involvement in pre-rRNA modification, processing, and folding remains unclear to what extent box C/D snoRNAs contribute to regulating the homeostasis of other cellular RNAs, including mRNAs. The data on interaction sites reported here may aid the elucidation of non-conventional roles of box C/D snoRNAs and the potential links between altered expression and cell differentiation, embryogenesis or human disease.

## ACKNOWLEDGEMENTS

We thank Aleksandra Helwak for critical reading of the MS. DT was supported by Wellcome [077248], TD and H.D-D were supported by BBSRC funding [Bb/L020416/1]. Work in the Wellcome Centre for Cell Biology is supported by core funding [092076]. DLJL was supported by the F.R.S./FNRS, the EJP-RD RiboEurope and DBAGenCure, the Université Libre de Bruxelles, and Région Wallonne (RiboCancer).

## DATA ACCESS

All sequence data from this study have been submitted to the NCBI Gene Expression Omnibus (GEO).

www.ncbi.nlm.nih.gov/geo/query/acc.cgi?acc=GSE114825

www.ncbi.nlm.nih.gov/geo/query/acc.cgi?acc=GSE121414

www.ncbi.nlm.nih.gov/geo/query/acc.cgi?acc=GSE121415

## SUPPLEMENTARY MATERIALS

**Dataset 1: Interactions recovered**

### Supplementary Figures

**Figure S1.**
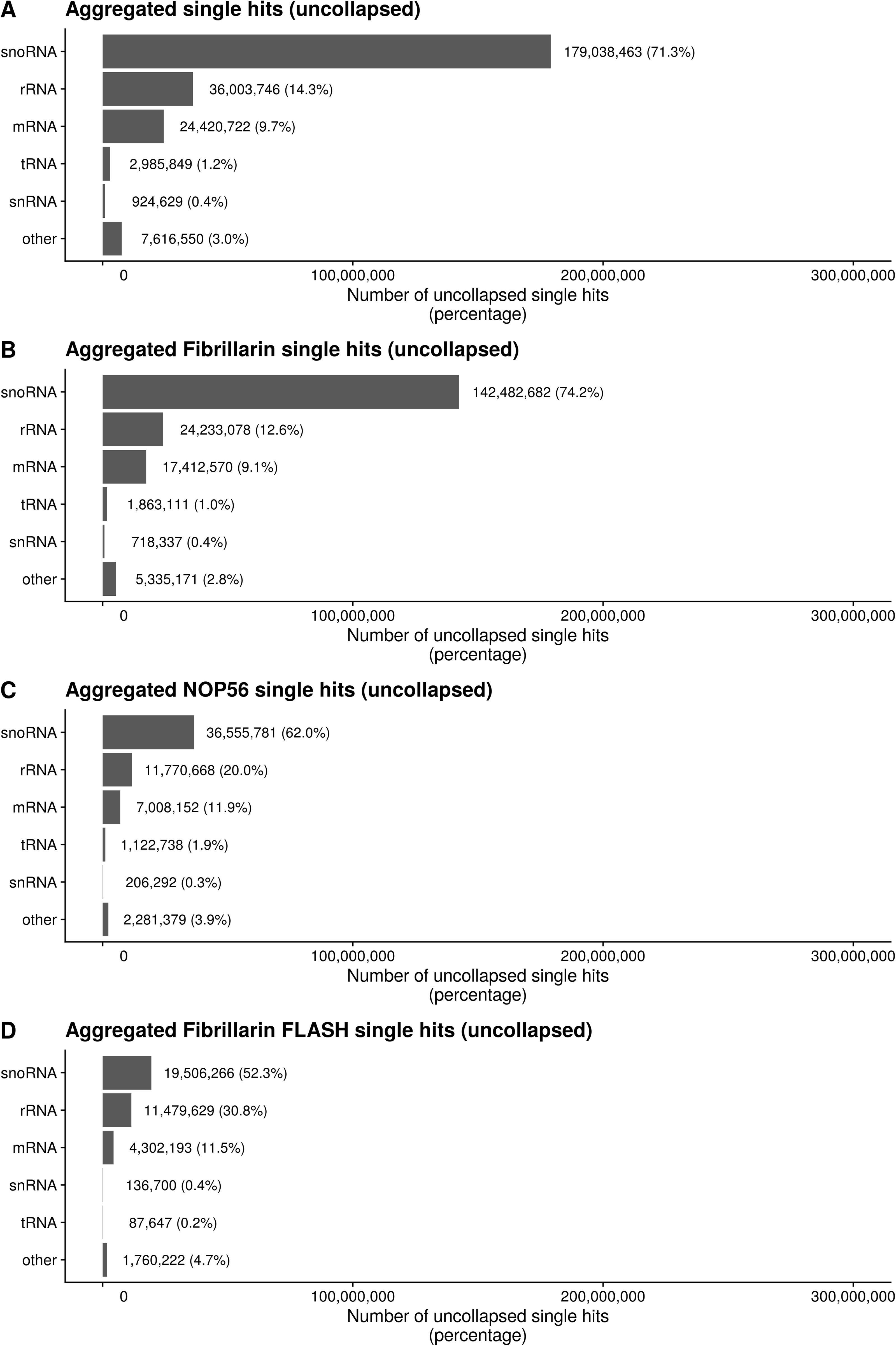
Single hits by biotype. (A) Breakdown of total uncollapsed single hit counts for different RNA biotypes. (B) Breakdown of uncollapsed single hit counts for different RNA biotypes in Fibrillarin CLASH and FLASH experiments. (C) Breakdown of uncollapsed single hit counts for different RNA biotypes in NOP56 CLASH and FLASH experiments. (D) Breakdown of uncollapsed single hit counts for different RNA biotypes in Fibrillarin FLASH experiments.

**Figure S2.**
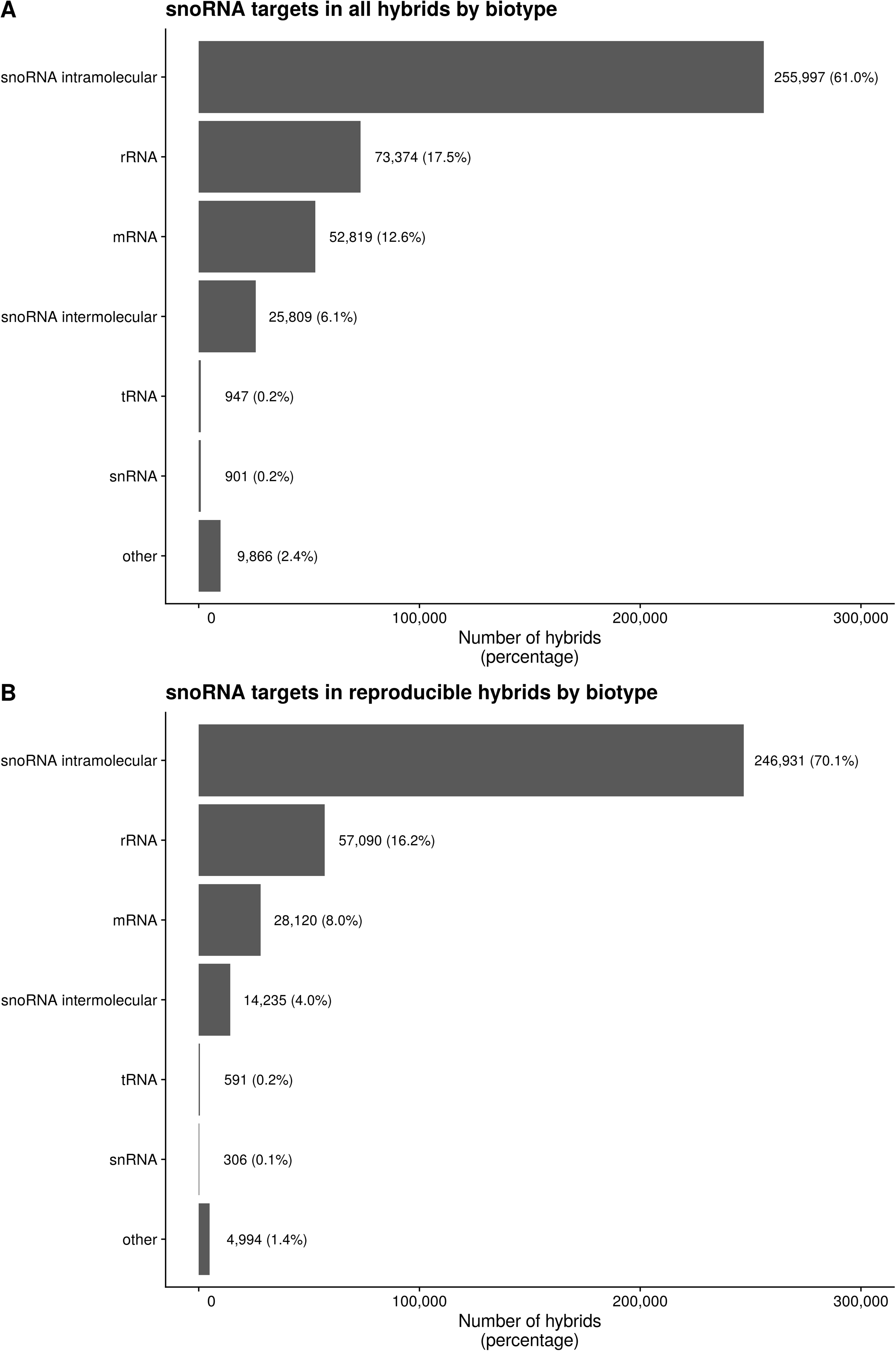
Hybrids recovered by biotype. (A) Breakdown of hybrid counts in snoRNA hybrids by target RNA biotype. (B) Breakdown of counts of reproducible, stable snoRNA hybrids (with predicted folding energy of -12dG or below) by target RNA biotype.

**Figure S3.**
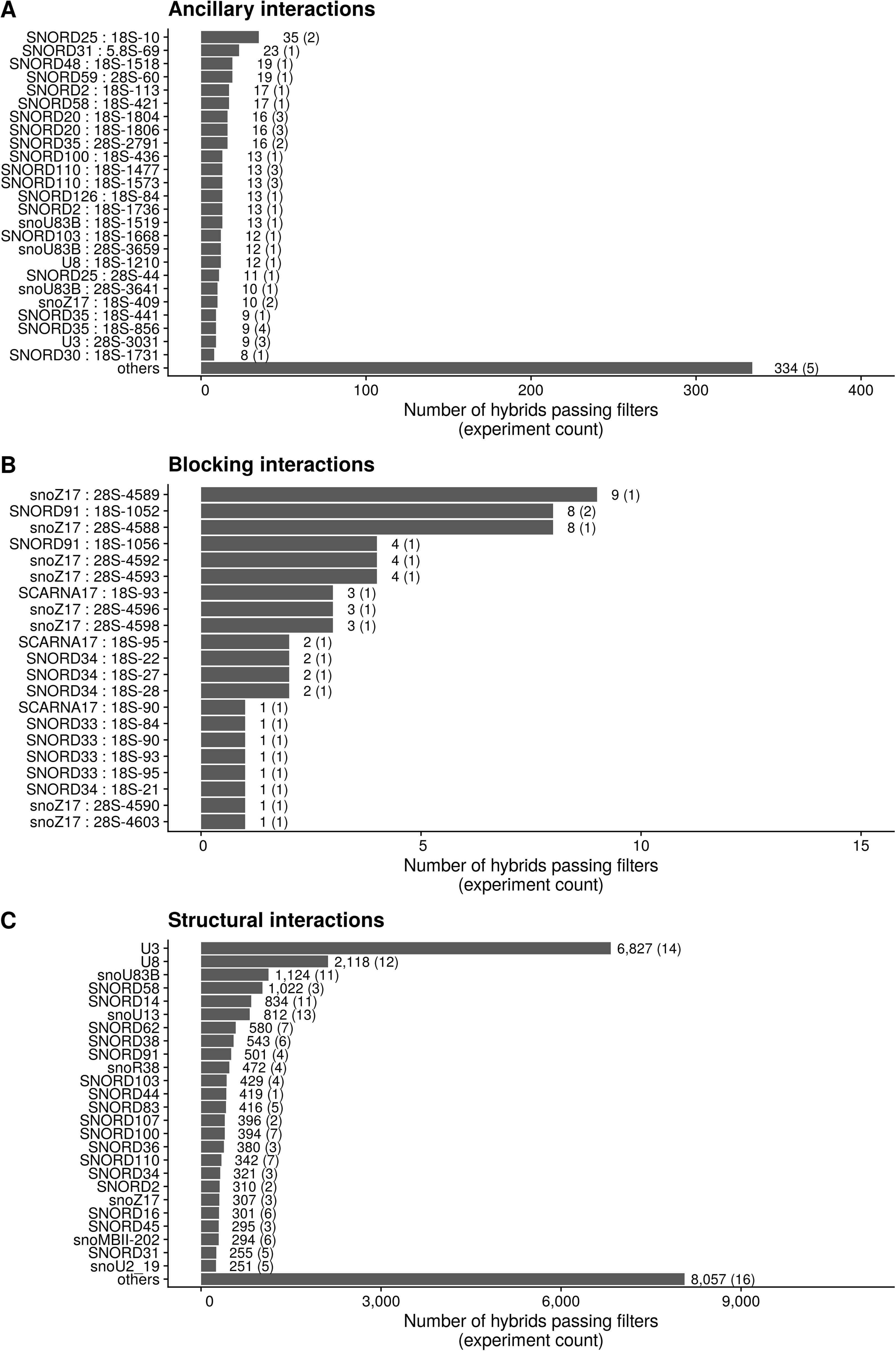
Classes of non-methylating rRNA interactions. (A, B) Breakdown of snoRNA-pre47S hybrids involved in ancillary interactions and blocking interactions respectively. The x axis labels show the snoRNA families involved, along with the co-ordinate of the relevant methylation site on the pre47S sequence. (C) Breakdown of snoRNA- pre47S hybrids involved in structural interactions. The x axis labels show the relevant snoRNA families.

**Figure S4.**
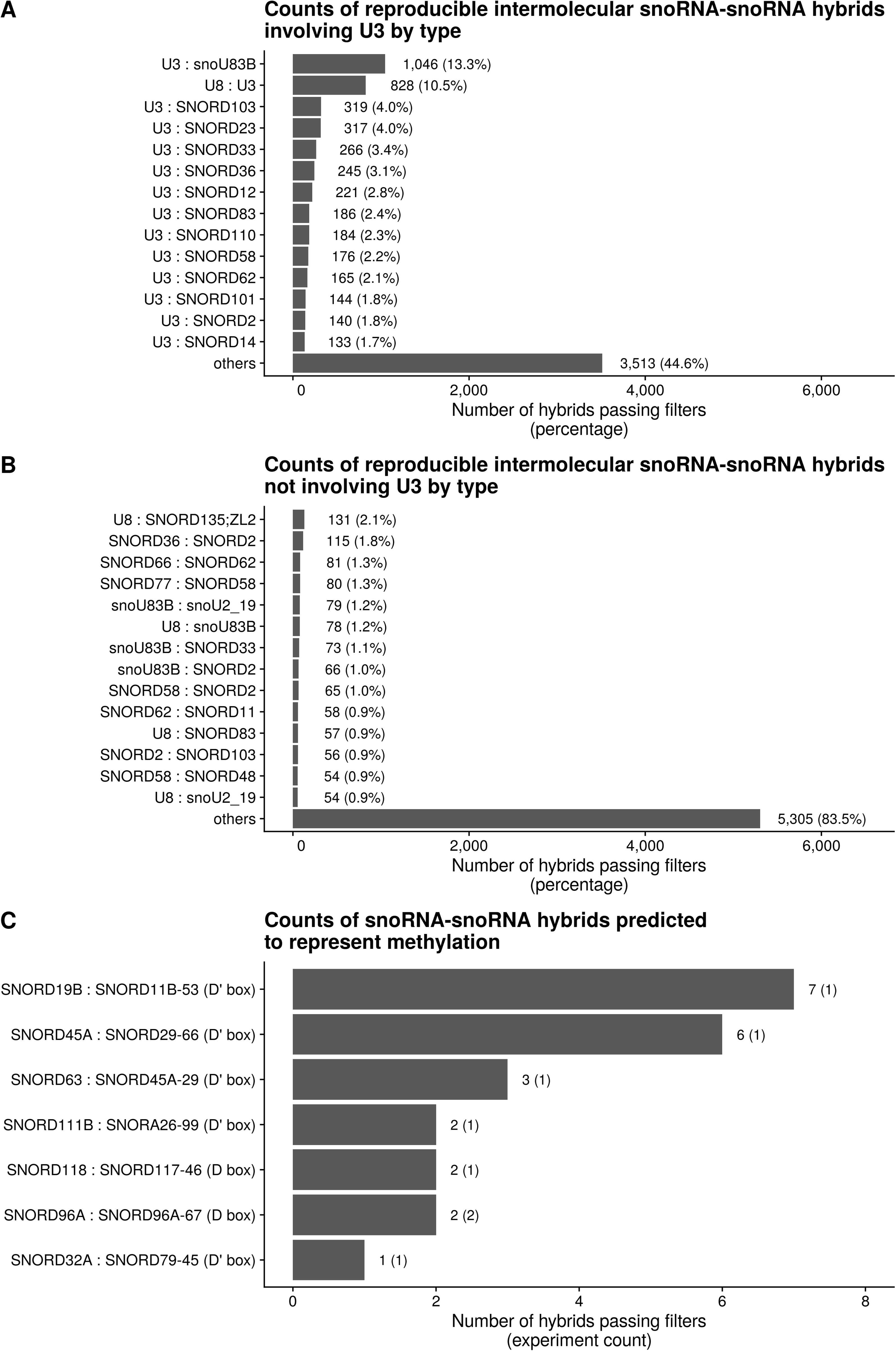
Characterization of snoRNA-snoRNA interactions. (A) Breakdown of reproducible intermolecular snoRNA-snoRNA hybrids by hybrid type. The hybrid type is obtained by joining the family name of each of the interacting snoRNAs. The order of the snoRNAs is not considered. (B) Breakdown of snoRNA-snoRNA hybrids that pass the filtering criteria described in figure 2A. The x axis labels show the snoRNAs and co-ordinate at which methylation is predicted to occur, and the snoRNA box 5 base pairs downstream of the methylating nucleotide. (C) Counts of snoRNA-snoRNA hybrids predicted to guide methylation with high confidence.

**Figure S5.**
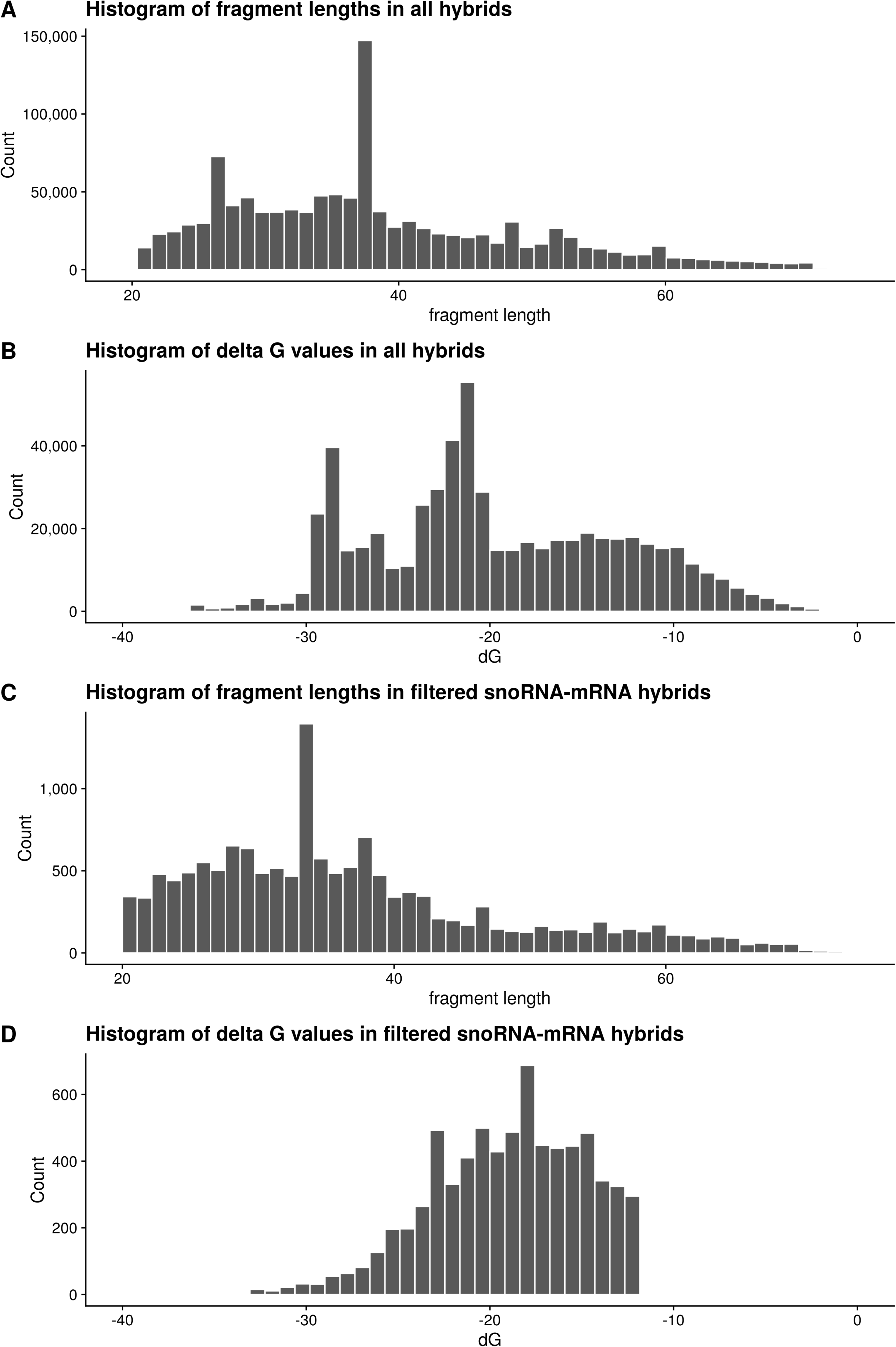
Distribution of length and stability of snoRNA-mRNA hybrids. (A) Histogram of fragment lengths in all hybrids. (B) Histogram of predicted folding energies in all hybrids. (C) Histogram of fragment lengths in reproducible, stable filtered snoRNA-mRNA hybrids with U3 and rRNA filters applied. (D) Histogram of predicted folding energies in reproducible, stable snoRNA-mRNA hybrids with U3 and rRNA filters applied.

**Figure S6.**
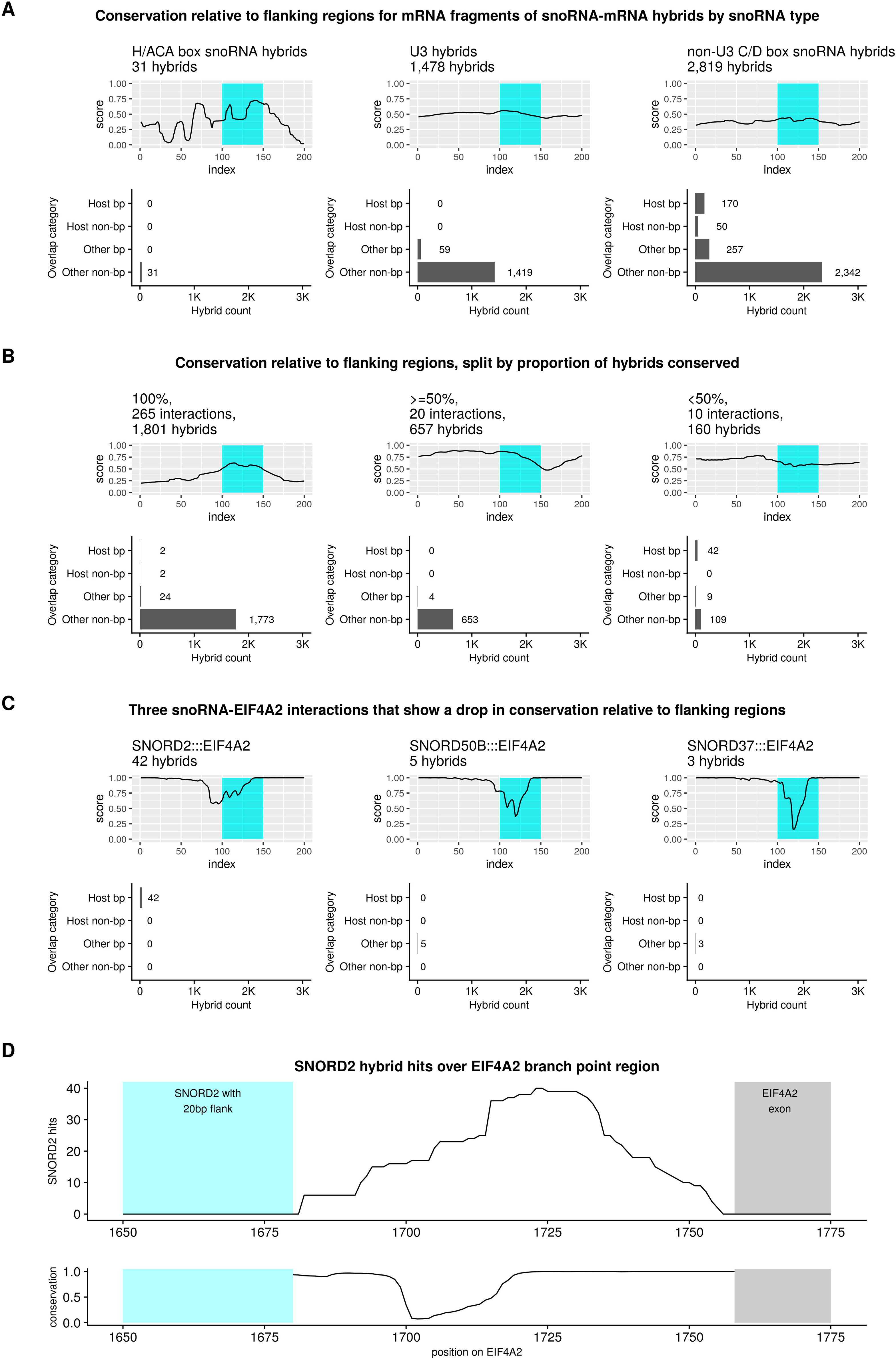
Conservation scores of snoRNA-mRNA interactions in human cells. (A) Conservation relative to flanking regions for mRNA fragments of snoRNA-mRNA hybrids involving U3, and non-U3 C/D box snoRNAs, respectively. Beneath each conservation plot is a bar chart showing the distribution of the mRNA fragments of the relevant hybrids between branch point regions of the host gene (host bp), non-branch point regions of the host gene (host non-bp), branch point regions of other genes (other bp) and non-branch point regions of other genes (other non-bp). (B) Conservation relative to flanking regions, split by proportion of hybrids conserved. These plots show that for the majority of interactions and hybrids, there is a clear peak in conservation corresponding to the location of the mRNA fragment of the hybrid. However, there is a small number of interactions with many hybrids that show a large decrease in conservation relative to flanks. This obscures the general pattern of conservation in the overall chart. (C) Three snoRNA-EIF4A2 interactions that show a drop in conservation relative to flanking regions. The three most common interactions with reduced conservation each involves an intron branch point region in EIF4A2. (D) Genome browser track showing the location of the mRNA fragment of SNORD2-EIF4A2 hybrids. In this case, the upstream flanking region overlaps the SNORD2 gene, and the downstream flanking region overlaps an exon of EIF4A2, both of which are highly conserved relative to the intron branch point region bound by the mRNA fragment of the hybrids, explaining the dip in conservation.

**Figure S7.**
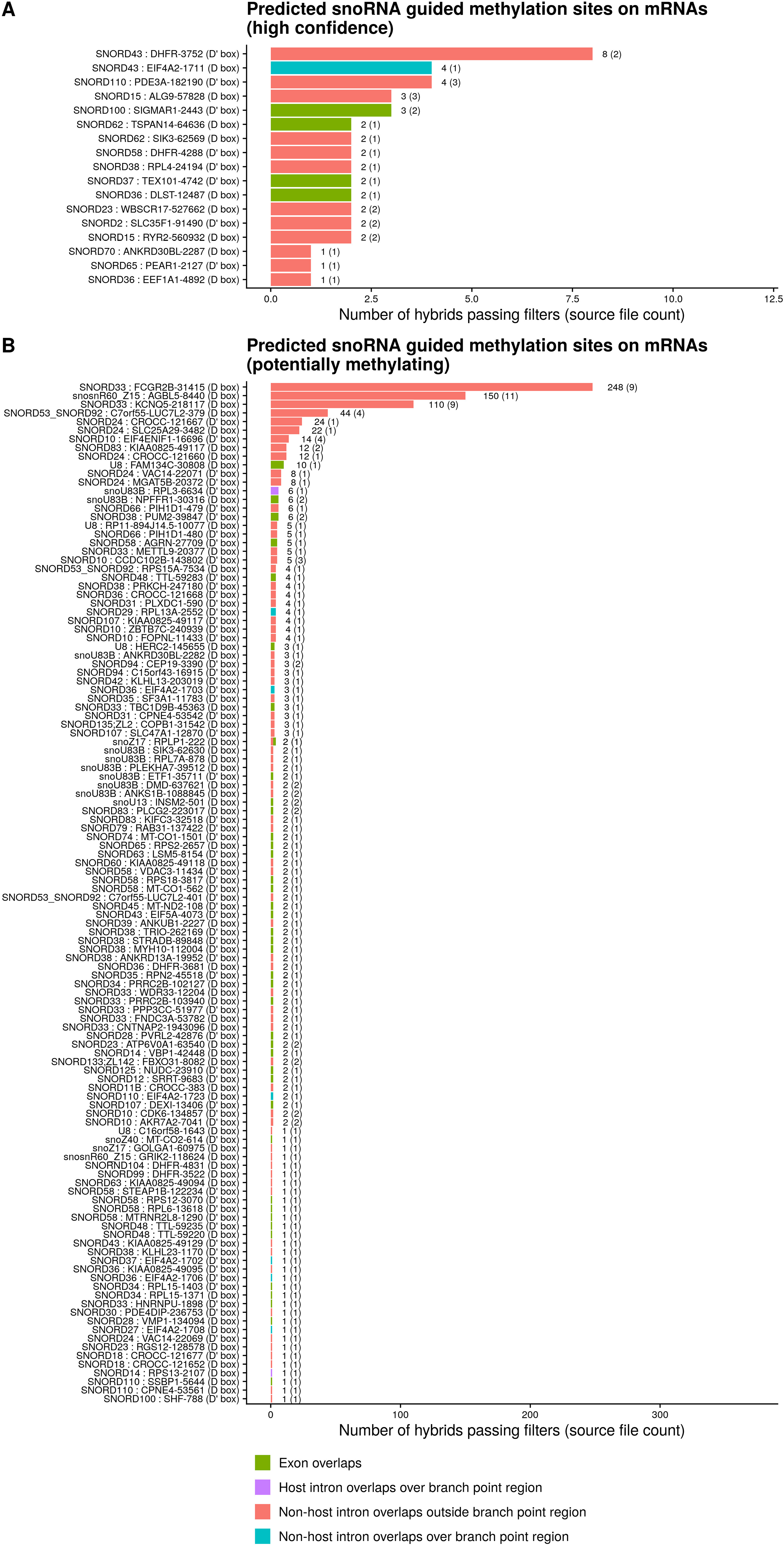
Potential methylating interactions with mRNAs. (A) Counts of filtered snoRNA-mRNA hybrids meeting the criteria for classification as high confidence methylating hybrids, broken down by interaction. (B) Counts of filtered snoRNA-mRNA hybrids meeting the criteria for classification as potentially methylating hybrids, broken down by interaction.

**Figure S8.**
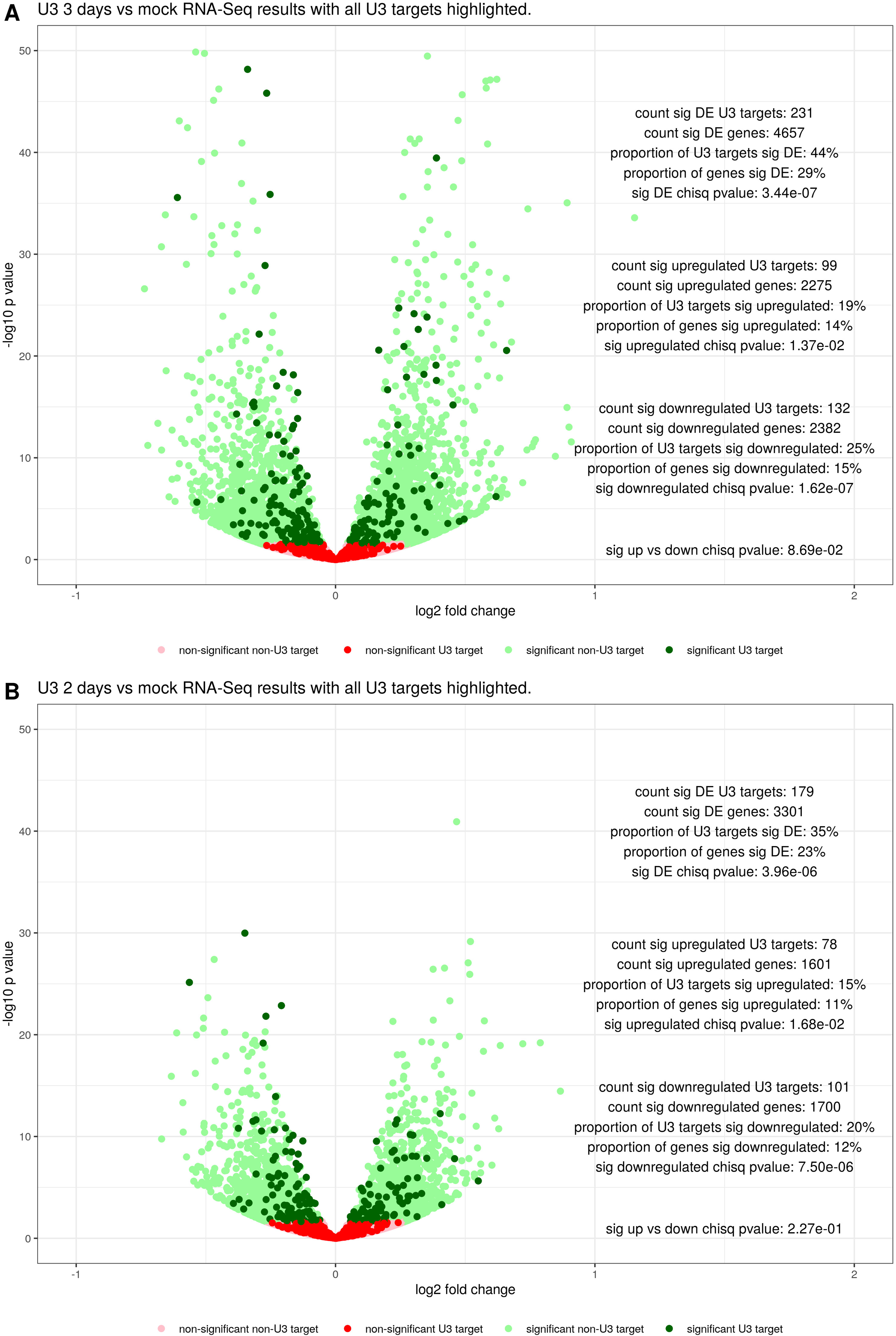
Changes in mRNA abundance following U3 depletion. (A) Volcano plot showing the changes in mRNA abundance following 3 days of U3 depletion, compared to a mock control, with U3 targets found in filtered U3-mRNA hybrids highlighted. (B) Volcano plot showing the changes in mRNA abundance following 2 days of U3 depletion, compared to a mock control, with U3 targets found in filtered U3-mRNA hybrids highlighted.

**Figure S9.** Filtering steps applied to hybrids. (A) A schematic representation of the filtering steps applied to different classes of RNA-RNA hybrids. Initial, single hit filtering was applied to all hybrids to filter out miscalled single reads. Filtering for reproducibility and stability was then applied to all hybrids. Additional U3 filtering was applied to all hybrids between snoRNAs and other biotypes to filter out miscalled U3-U3 intermolecular hybrids. Finally, additional rRNA filtering was applied to all hybrids between snoRNAs and other non-rRNA biotypes, to filter out miscalled snoRNA-rRNA hybrids. (B) Profile plot showing the distribution of filtered, reproducible SNORD3A (U3) hybrids over SNORD3A by biotype. Each line shows the profile for a different SNORD3A target biotype. The grey shaded regions represent 20 base pair flanking regions upstream and downstream of SNORD3A. The light blue shaded region represents the D box of SNORD3A, the purple line shows the nucleotide 5 base pairs upstream of the D box of SNORD3A, and the light green shaded region represents the D’ box of SNORD3A.

### Supplementary tables

**Table S1:**
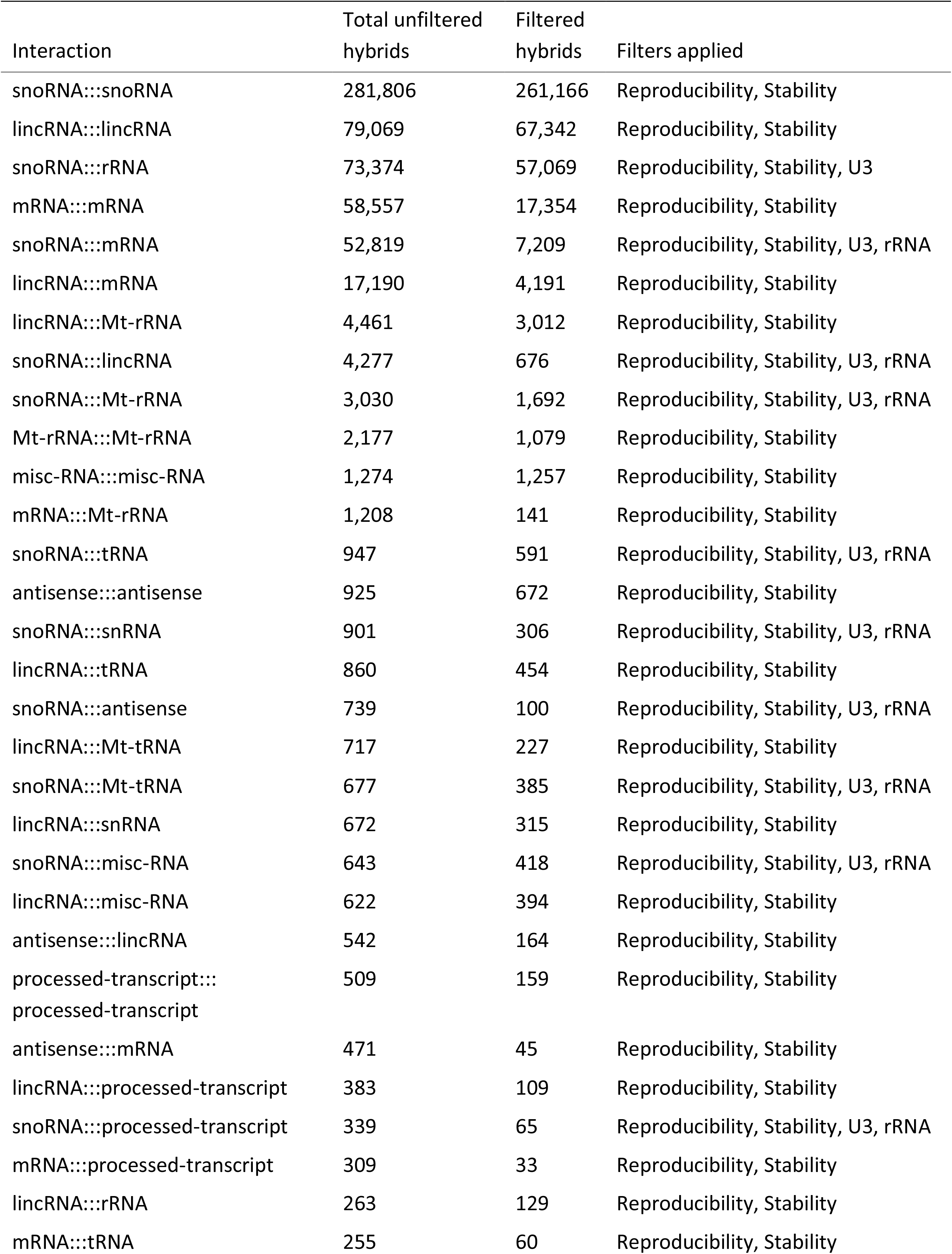
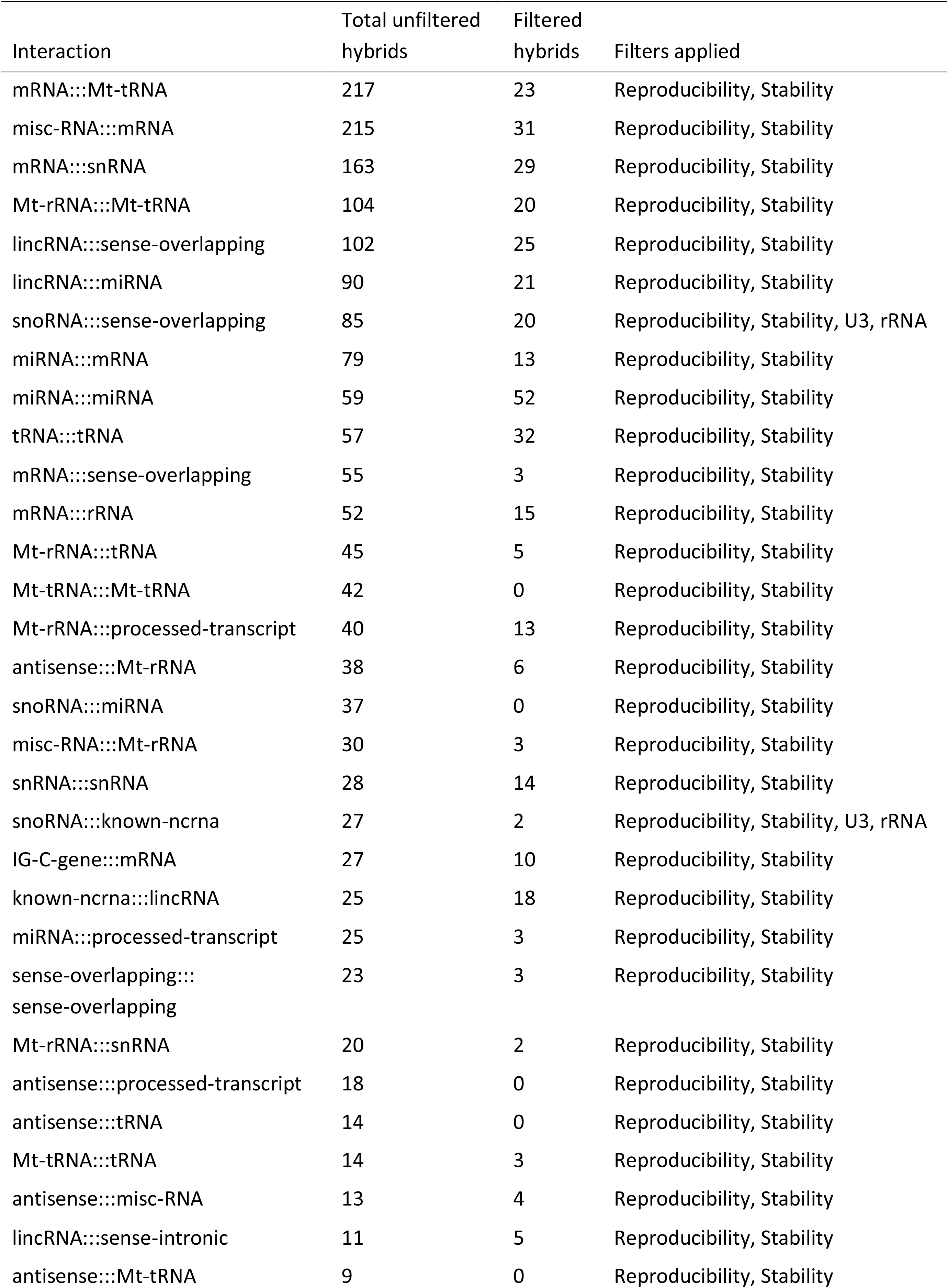
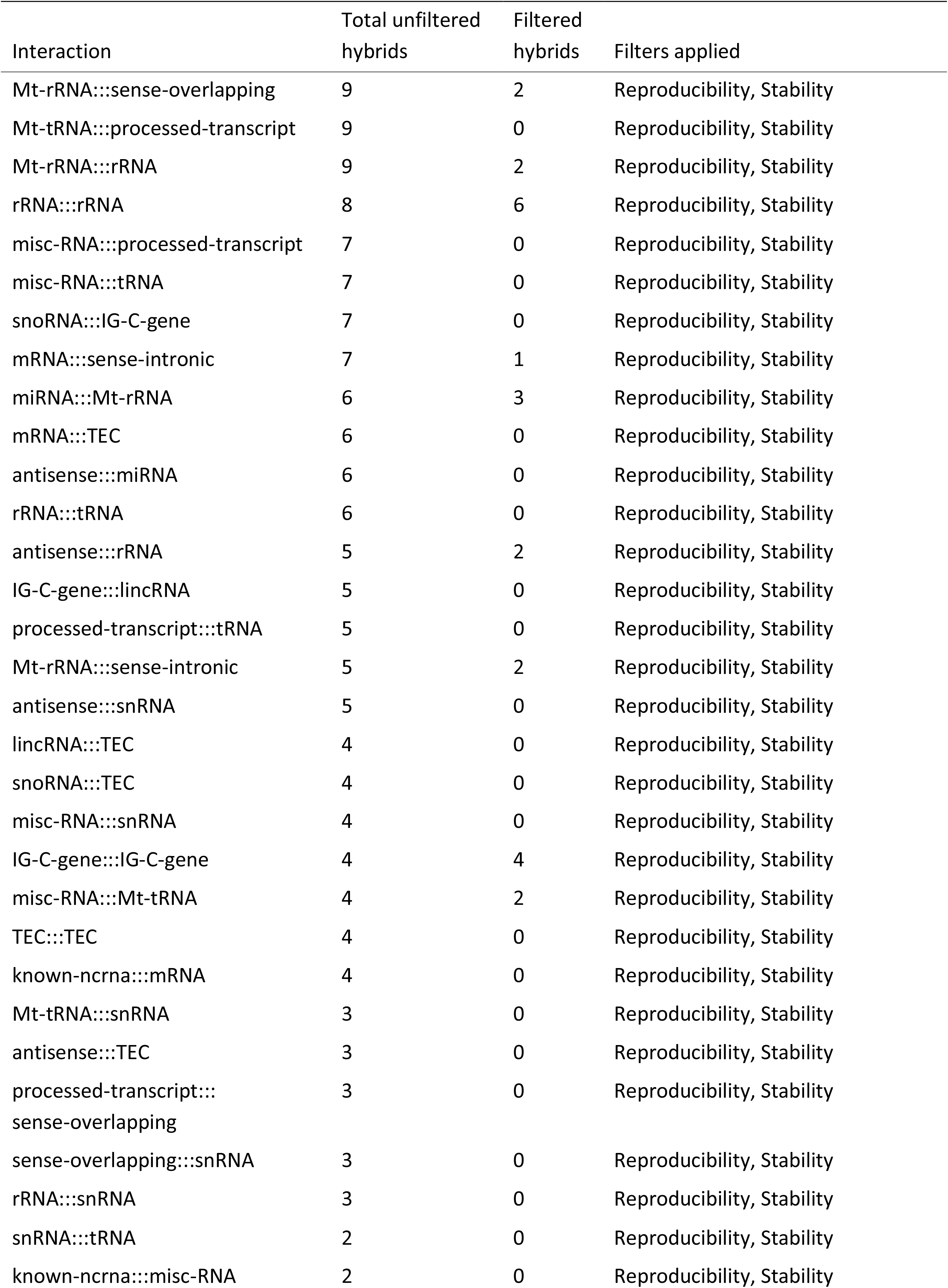
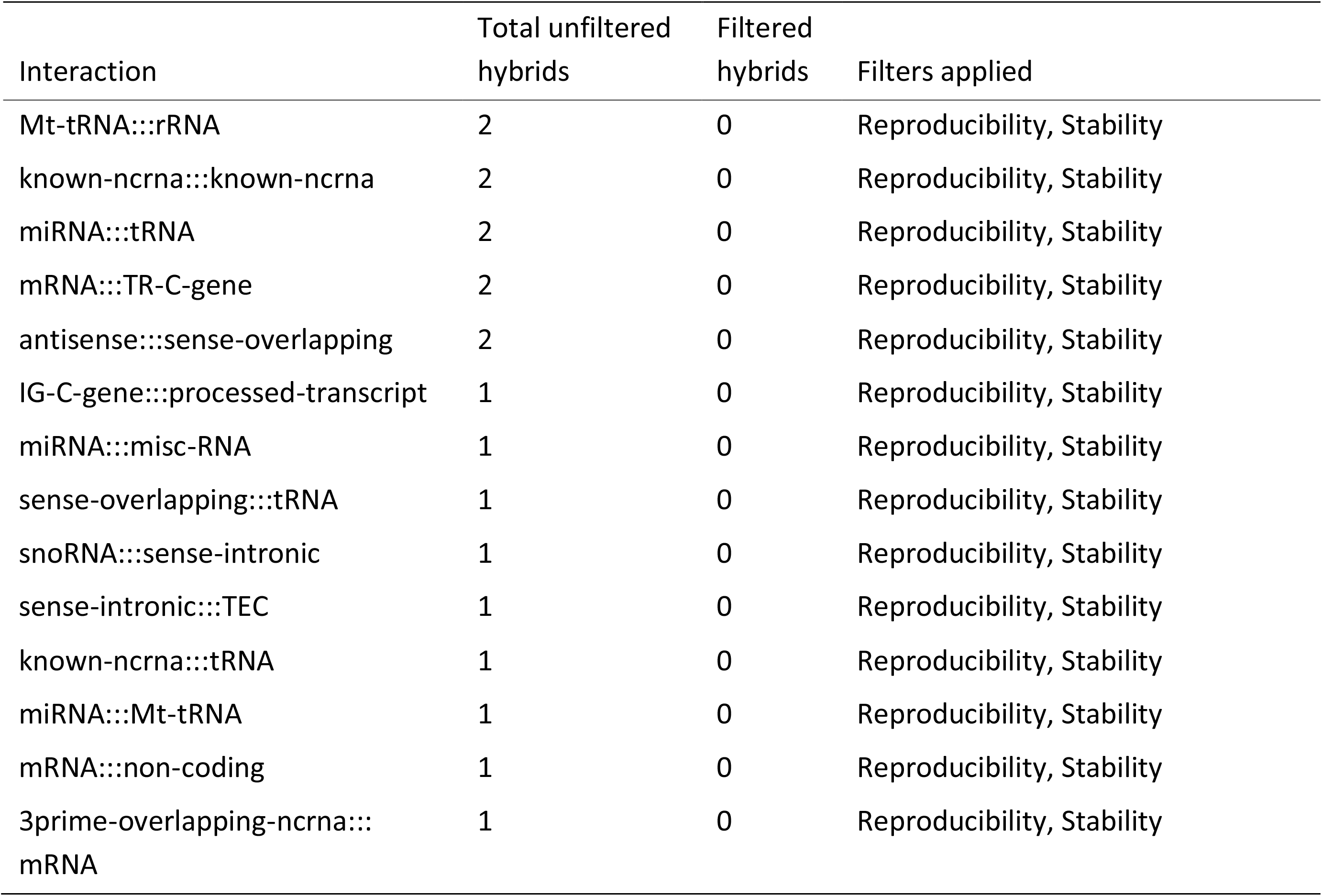
Overall hybrid counts by biotype.

| Interaction | Total unfiltered hybrids | Filtered hybrids | Filters applied |
| --- | --- | --- | --- |
| snoRNA:::snoRNA | 281,806 | 261,166 | Reproducibility, Stability |
| lincRNA:::lincRNA | 79,069 | 67,342 | Reproducibility, Stability |
| snoRNA:::rRNA | 73,374 | 57,069 | Reproducibility, Stability, U3 |
| mRNA:::mRNA | 58,557 | 17,354 | Reproducibility, Stability |
| snoRNA:::mRNA | 52,819 | 7,209 | Reproducibility, Stability, U3, rRNA |
| lincRNA:::mRNA | 17,190 | 4,191 | Reproducibility, Stability |
| lincRNA:::Mt-rRNA | 4,461 | 3,012 | Reproducibility, Stability |
| snoRNA:::lincRNA | 4,277 | 676 | Reproducibility, Stability, U3, rRNA |
| snoRNA:::Mt-rRNA | 3,030 | 1,692 | Reproducibility, Stability, U3, rRNA |
| Mt-rRNA:::Mt-rRNA | 2,177 | 1,079 | Reproducibility, Stability |
| misc-RNA:::misc-RNA | 1,274 | 1,257 | Reproducibility, Stability |
| mRNA:::Mt-rRNA | 1,208 | 141 | Reproducibility, Stability |
| snoRNA:::tRNA | 947 | 591 | Reproducibility, Stability, U3, rRNA |
| antisense:::antisense | 925 | 672 | Reproducibility, Stability |
| snoRNA:::snRNA | 901 | 306 | Reproducibility, Stability, U3, rRNA |
| lincRNA:::tRNA | 860 | 454 | Reproducibility, Stability |
| snoRNA:::antisense | 739 | 100 | Reproducibility, Stability, U3, rRNA |
| lincRNA:::Mt-tRNA | 717 | 227 | Reproducibility, Stability |
| snoRNA:::Mt-tRNA | 677 | 385 | Reproducibility, Stability, U3, rRNA |
| lincRNA:::snRNA | 672 | 315 | Reproducibility, Stability |
| snoRNA:::misc-RNA | 643 | 418 | Reproducibility, Stability, U3, rRNA |
| lincRNA:::misc-RNA | 622 | 394 | Reproducibility, Stability |
| antisense:::lincRNA | 542 | 164 | Reproducibility, Stability |
| processed-transcript:::<br>processed-transcript | 509 | 159 | Reproducibility, Stability |
| antisense:::mRNA | 471 | 45 | Reproducibility, Stability |
| lincRNA:::processed-transcript | 383 | 109 | Reproducibility, Stability |
| snoRNA:::processed-transcript | 339 | 65 | Reproducibility, Stability, U3, rRNA |
| mRNA:::processed-transcript | 309 | 33 | Reproducibility, Stability |
| lincRNA:::rRNA | 263 | 129 | Reproducibility, Stability |
| mRNA:::tRNA | 255 | 60 | Reproducibility, Stability |
| mRNA:::Mt-tRNA | 217 | 23 | Reproducibility, Stability |
| misc-RNA:::mRNA | 215 | 31 | Reproducibility, Stability |
| mRNA:::snRNA | 163 | 29 | Reproducibility, Stability |
| Mt-rRNA:::Mt-tRNA | 104 | 20 | Reproducibility, Stability |
| lincRNA:::sense-overlapping | 102 | 25 | Reproducibility, Stability |
| lincRNA:::miRNA | 90 | 21 | Reproducibility, Stability |
| snoRNA:::sense-overlapping | 85 | 20 | Reproducibility, Stability, U3, rRNA |
| miRNA:::mRNA | 79 | 13 | Reproducibility, Stability |
| miRNA:::miRNA | 59 | 52 | Reproducibility, Stability |
| tRNA:::tRNA | 57 | 32 | Reproducibility, Stability |
| mRNA:::sense-overlapping | 55 | 3 | Reproducibility, Stability |
| mRNA:::rRNA | 52 | 15 | Reproducibility, Stability |
| Mt-rRNA:::tRNA | 45 | 5 | Reproducibility, Stability |
| Mt-tRNA:::Mt-tRNA | 42 | 0 | Reproducibility, Stability |
| Mt-rRNA:::processed-transcript | 40 | 13 | Reproducibility, Stability |
| antisense:::Mt-rRNA | 38 | 6 | Reproducibility, Stability |
| snoRNA:::miRNA | 37 | 0 | Reproducibility, Stability |
| misc-RNA:::Mt-rRNA | 30 | 3 | Reproducibility, Stability |
| snRNA:::snRNA | 28 | 14 | Reproducibility, Stability |
| snoRNA:::known-ncrna | 27 | 2 | Reproducibility, Stability, U3, rRNA |
| IG-C-gene:::mRNA | 27 | 10 | Reproducibility, Stability |
| known-ncrna:::lincRNA | 25 | 18 | Reproducibility, Stability |
| miRNA:::processed-transcript | 25 | 3 | Reproducibility, Stability |
| sense-overlapping:::<br>sense-overlapping | 23 | 3 | Reproducibility, Stability |
| Mt-rRNA:::snRNA | 20 | 2 | Reproducibility, Stability |
| antisense:::processed-transcript | 18 | 0 | Reproducibility, Stability |
| antisense:::tRNA | 14 | 0 | Reproducibility, Stability |
| Mt-tRNA:::tRNA | 14 | 3 | Reproducibility, Stability |
| antisense:::misc-RNA | 13 | 4 | Reproducibility, Stability |
| lincRNA:::sense-intronic | 11 | 5 | Reproducibility, Stability |
| antisense:::Mt-tRNA | 9 | 0 | Reproducibility, Stability |
| Mt-rRNA:::sense-overlapping | 9 | 2 | Reproducibility, Stability |
| Mt-tRNA:::processed-transcript | 9 | 0 | Reproducibility, Stability |
| Mt-rRNA:::rRNA | 9 | 2 | Reproducibility, Stability |
| rRNA:::rRNA | 8 | 6 | Reproducibility, Stability |
| misc-RNA:::processed-transcript | 7 | 0 | Reproducibility, Stability |
| misc-RNA:::tRNA | 7 | 0 | Reproducibility, Stability |
| snoRNA:::IG-C-gene | 7 | 0 | Reproducibility, Stability |
| mRNA:::sense-intronic | 7 | 1 | Reproducibility, Stability |
| miRNA:::Mt-rRNA | 6 | 3 | Reproducibility, Stability |
| mRNA:::TEC | 6 | 0 | Reproducibility, Stability |
| antisense:::miRNA | 6 | 0 | Reproducibility, Stability |
| rRNA:::tRNA | 6 | 0 | Reproducibility, Stability |
| antisense:::rRNA | 5 | 2 | Reproducibility, Stability |
| IG-C-gene:::lincRNA | 5 | 0 | Reproducibility, Stability |
| processed-transcript:::tRNA | 5 | 0 | Reproducibility, Stability |
| Mt-rRNA:::sense-intronic | 5 | 2 | Reproducibility, Stability |
| antisense:::snRNA | 5 | 0 | Reproducibility, Stability |
| lincRNA:::TEC | 4 | 0 | Reproducibility, Stability |
| snoRNA:::TEC | 4 | 0 | Reproducibility, Stability |
| misc-RNA:::snRNA | 4 | 0 | Reproducibility, Stability |
| IG-C-gene:::IG-C-gene | 4 | 4 | Reproducibility, Stability |
| misc-RNA:::Mt-tRNA | 4 | 2 | Reproducibility, Stability |
| TEC:::TEC | 4 | 0 | Reproducibility, Stability |
| known-ncrna:::mRNA | 4 | 0 | Reproducibility, Stability |
| Mt-tRNA:::snRNA | 3 | 0 | Reproducibility, Stability |
| antisense:::TEC | 3 | 0 | Reproducibility, Stability |
| processed-transcript:::<br>sense-overlapping | 3 | 0 | Reproducibility, Stability |
| sense-overlapping:::snRNA | 3 | 0 | Reproducibility, Stability |
| rRNA:::snRNA | 3 | 0 | Reproducibility, Stability |
| snRNA:::tRNA | 2 | 0 | Reproducibility, Stability |
| known-ncrna:::misc-RNA | 2 | 0 | Reproducibility, Stability |
| Mt-tRNA:::rRNA | 2 | 0 | Reproducibility, Stability |
| known-ncrna:::known-ncrna | 2 | 0 | Reproducibility, Stability |
| miRNA:::tRNA | 2 | 0 | Reproducibility, Stability |
| mRNA:::TR-C-gene | 2 | 0 | Reproducibility, Stability |
| antisense:::sense-overlapping | 2 | 0 | Reproducibility, Stability |
| IG-C-gene:::processed-transcript | 1 | 0 | Reproducibility, Stability |
| miRNA:::misc-RNA | 1 | 0 | Reproducibility, Stability |
| sense-overlapping:::tRNA | 1 | 0 | Reproducibility, Stability |
| snoRNA:::sense-intronic | 1 | 0 | Reproducibility, Stability |
| sense-intronic:::TEC | 1 | 0 | Reproducibility, Stability |
| known-ncrna:::tRNA | 1 | 0 | Reproducibility, Stability |
| miRNA:::Mt-tRNA | 1 | 0 | Reproducibility, Stability |
| mRNA:::non-coding | 1 | 0 | Reproducibility, Stability |
| 3prime-overlapping-ncrna::: mRNA | 1 | 0 | Reproducibility, Stability |

**Table S2:**
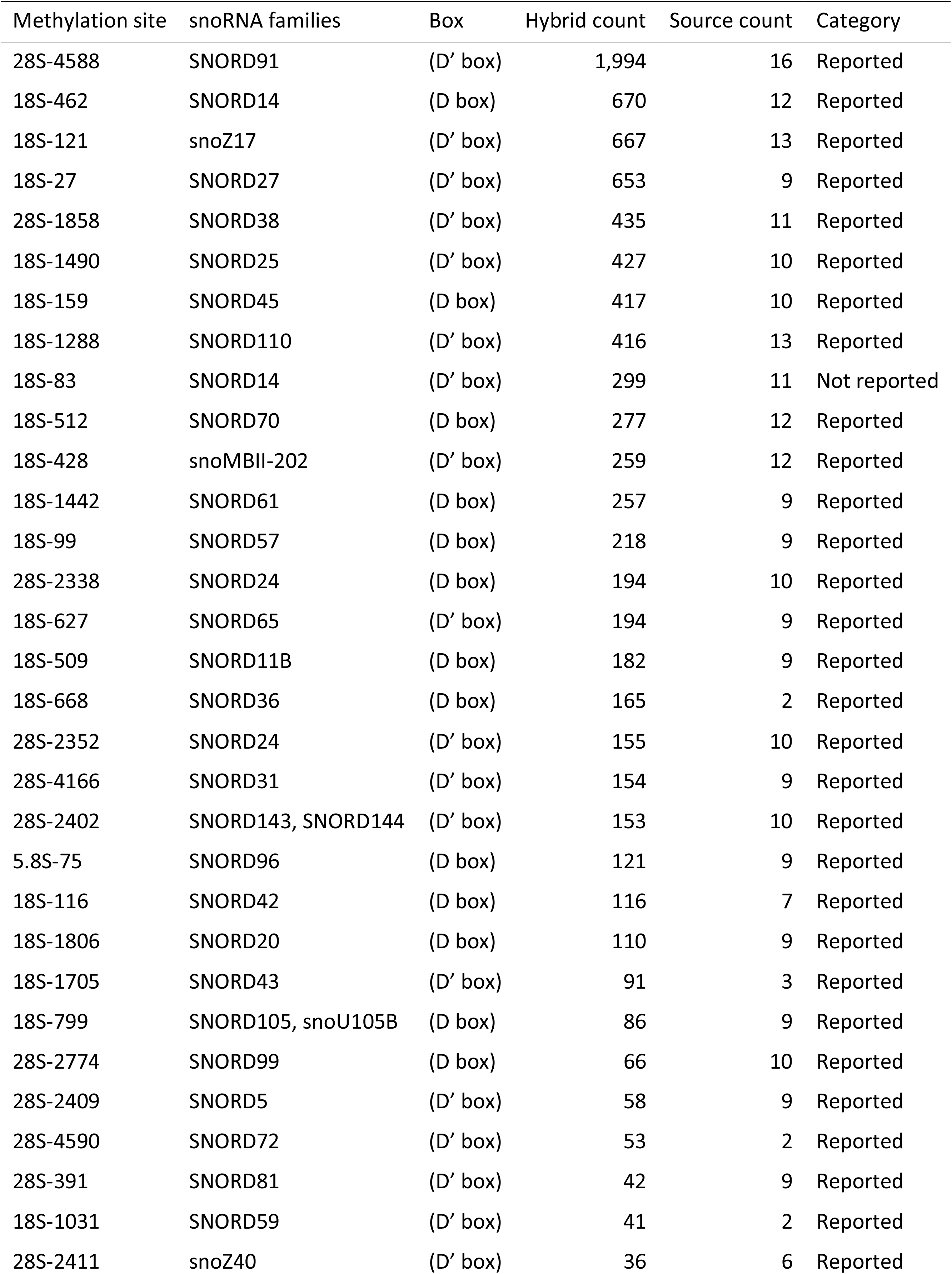
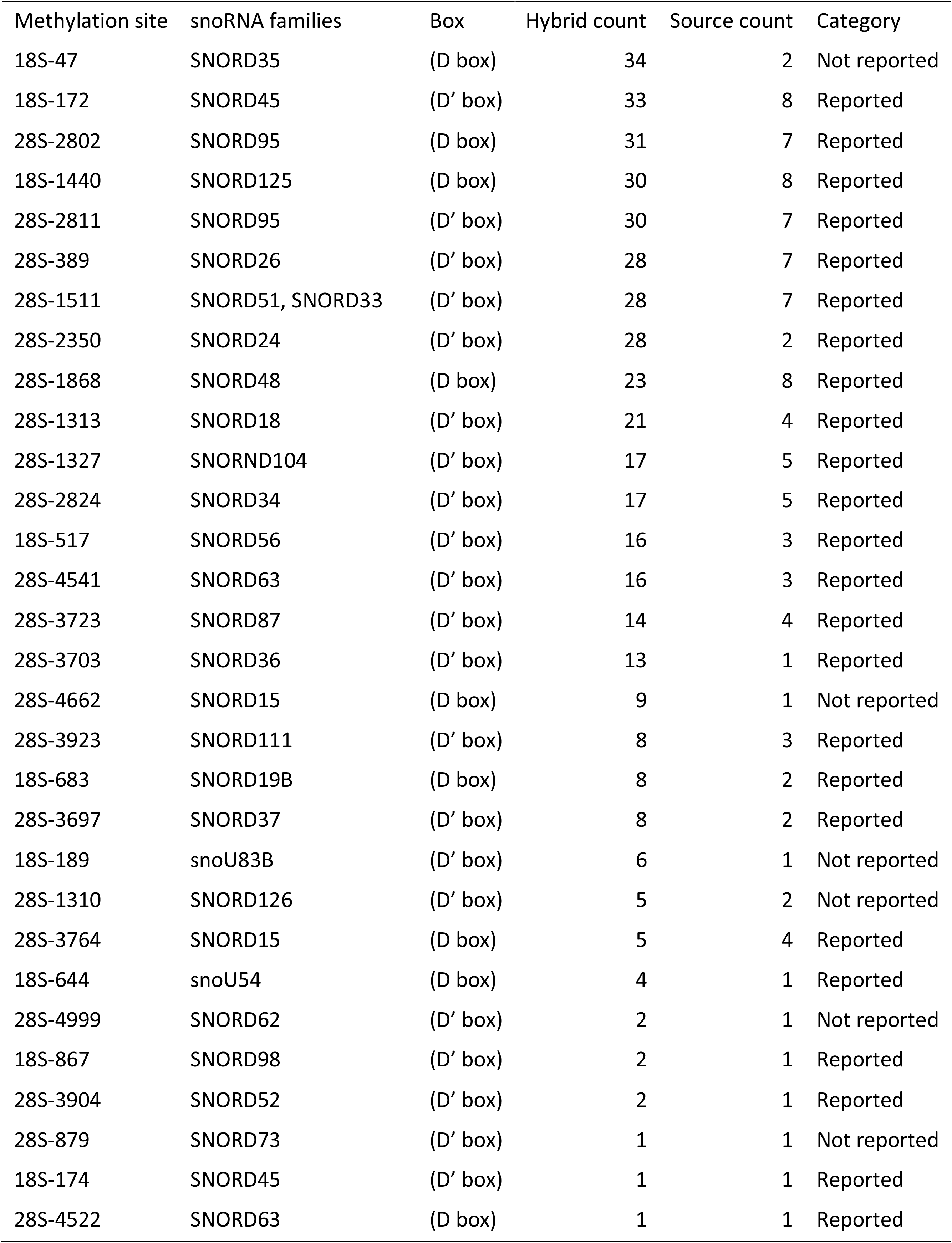
High confidence methylating hybrids.

| Methylation site | snoRNA families | Box | Hybrid count | Source count | Category |
| --- | --- | --- | --- | --- | --- |
| 28S-4588 | SNORD91 | (D' box) | 1,994 | 16 | Reported |
| 18S-462 | SNORD14 | (D box) | 670 | 12 | Reported |
| 18S-121 | snoZ17 | (D' box) | 667 | 13 | Reported |
| 18S-27 | SNORD27 | (D' box) | 653 | 9 | Reported |
| 28S-1858 | SNORD38 | (D' box) | 435 | 11 | Reported |
| 18S-1490 | SNORD25 | (D' box) | 427 | 10 | Reported |
| 18S-159 | SNORD45 | (D box) | 417 | 10 | Reported |
| 18S-1288 | SNORD110 | (D' box) | 416 | 13 | Reported |
| 18S-83 | SNORD14 | (D' box) | 299 | 11 | Not reported |
| 18S-512 | SNORD70 | (D box) | 277 | 12 | Reported |
| 18S-428 | snoMBII-202 | (D' box) | 259 | 12 | Reported |
| 18S-1442 | SNORD61 | (D box) | 257 | 9 | Reported |
| 18S-99 | SNORD57 | (D box) | 218 | 9 | Reported |
| 28S-2338 | SNORD24 | (D box) | 194 | 10 | Reported |
| 18S-627 | SNORD65 | (D' box) | 194 | 9 | Reported |
| 18S-509 | SNORD11B | (D box) | 182 | 9 | Reported |
| 18S-668 | SNORD36 | (D box) | 165 | 2 | Reported |
| 28S-2352 | SNORD24 | (D' box) | 155 | 10 | Reported |
| 28S-4166 | SNORD31 | (D' box) | 154 | 9 | Reported |
| 28S-2402 | SNORD143, SNORD144 | (D' box) | 153 | 10 | Reported |
| 5.8S-75 | SNORD96 | (D box) | 121 | 9 | Reported |
| 18S-116 | SNORD42 | (D box) | 116 | 7 | Reported |
| 18S-1806 | SNORD20 | (D box) | 110 | 9 | Reported |
| 18S-1705 | SNORD43 | (D' box) | 91 | 3 | Reported |
| 18S-799 | SNORD105, snoU105B | (D box) | 86 | 9 | Reported |
| 28S-2774 | SNORD99 | (D box) | 66 | 10 | Reported |
| 28S-2409 | SNORD5 | (D' box) | 58 | 9 | Reported |
| 28S-4590 | SNORD72 | (D' box) | 53 | 2 | Reported |
| 28S-391 | SNORD81 | (D' box) | 42 | 9 | Reported |
| 18S-1031 | SNORD59 | (D' box) | 41 | 2 | Reported |
| 28S-2411 | snoZ40 | (D' box) | 36 | 6 | Reported |
| 18S-47 | SNORD35 | (D box) | 34 | 2 | Not reported |
| 18S-172 | SNORD45 | (D' box) | 33 | 8 | Reported |
| 28S-2802 | SNORD95 | (D box) | 31 | 7 | Reported |
| 18S-1440 | SNORD125 | (D box) | 30 | 8 | Reported |
| 28S-2811 | SNORD95 | (D' box) | 30 | 7 | Reported |
| 28S-389 | SNORD26 | (D' box) | 28 | 7 | Reported |
| 28S-1511 | SNORD51, SNORD33 | (D' box) | 28 | 7 | Reported |
| 28S-2350 | SNORD24 | (D' box) | 28 | 2 | Reported |
| 28S-1868 | SNORD48 | (D box) | 23 | 8 | Reported |
| 28S-1313 | SNORD18 | (D' box) | 21 | 4 | Reported |
| 28S-1327 | SNORND104 | (D' box) | 17 | 5 | Reported |
| 28S-2824 | SNORD34 | (D' box) | 17 | 5 | Reported |
| 18S-517 | SNORD56 | (D' box) | 16 | 3 | Reported |
| 28S-4541 | SNORD63 | (D' box) | 16 | 3 | Reported |
| 28S-3723 | SNORD87 | (D' box) | 14 | 4 | Reported |
| 28S-3703 | SNORD36 | (D' box) | 13 | 1 | Reported |
| 28S-4662 | SNORD15 | (D box) | 9 | 1 | Not reported |
| 28S-3923 | SNORD111 | (D' box) | 8 | 3 | Reported |
| 18S-683 | SNORD19B | (D box) | 8 | 2 | Reported |
| 28S-3697 | SNORD37 | (D' box) | 8 | 2 | Reported |
| 18S-189 | snoU83B | (D' box) | 6 | 1 | Not reported |
| 28S-1310 | SNORD126 | (D' box) | 5 | 2 | Not reported |
| 28S-3764 | SNORD15 | (D box) | 5 | 4 | Reported |
| 18S-644 | snoU54 | (D box) | 4 | 1 | Reported |
| 28S-4999 | SNORD62 | (D' box) | 2 | 1 | Not reported |
| 18S-867 | SNORD98 | (D' box) | 2 | 1 | Reported |
| 28S-3904 | SNORD52 | (D' box) | 2 | 1 | Reported |
| 28S-879 | SNORD73 | (D' box) | 1 | 1 | Not reported |
| 18S-174 | SNORD45 | (D' box) | 1 | 1 | Reported |
| 28S-4522 | SNORD63 | (D box) | 1 | 1 | Reported |

**Table S3:**
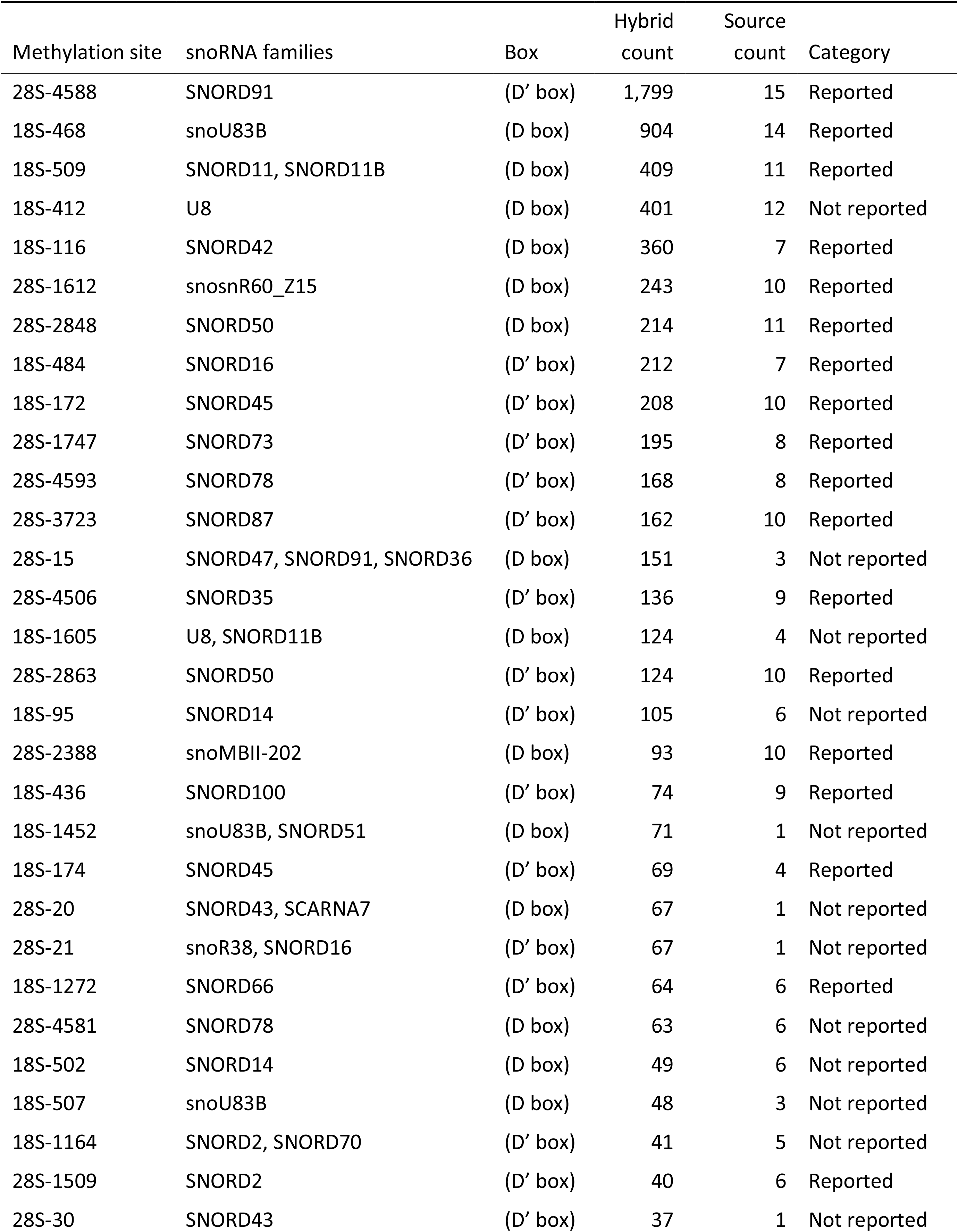
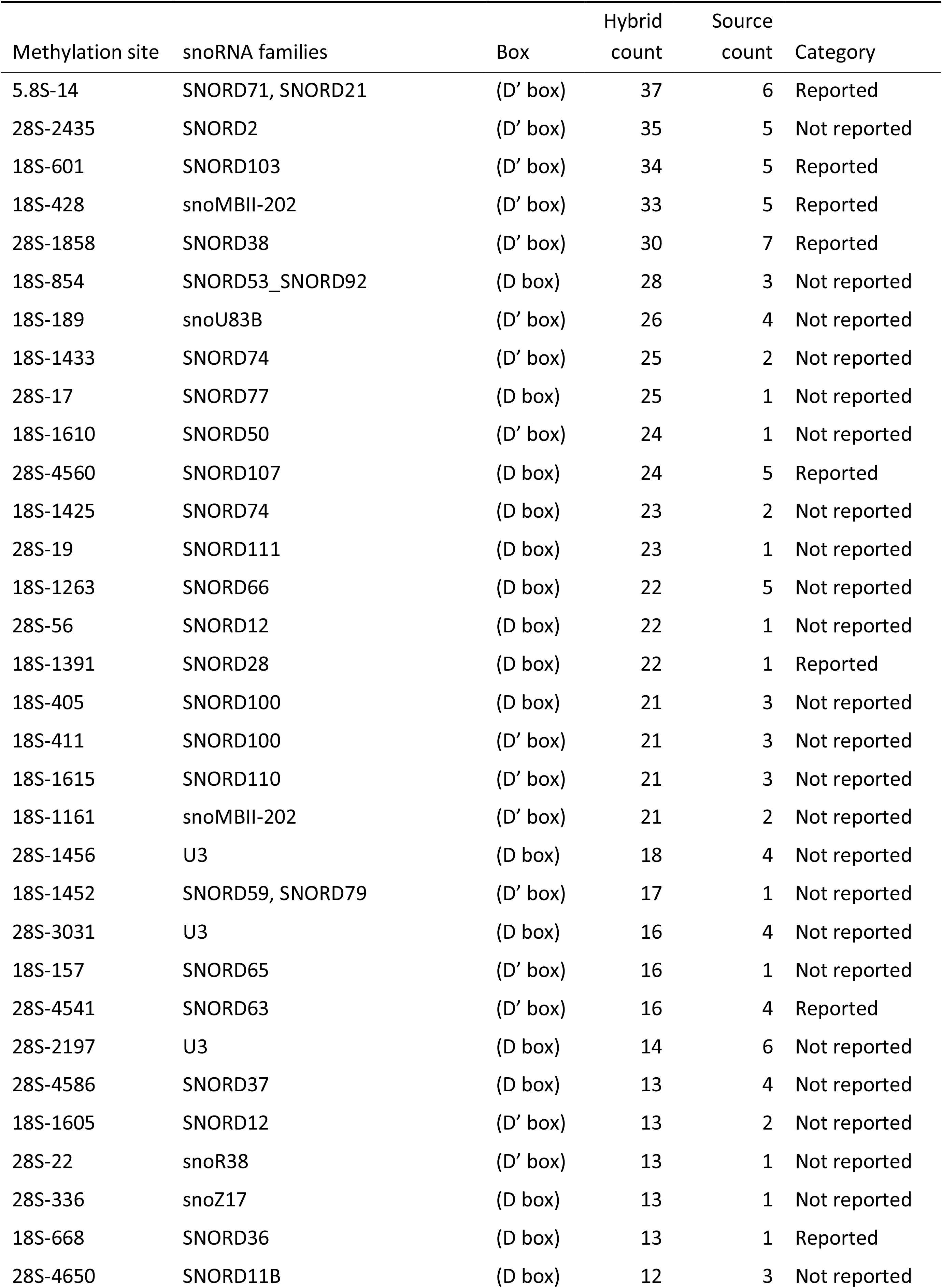
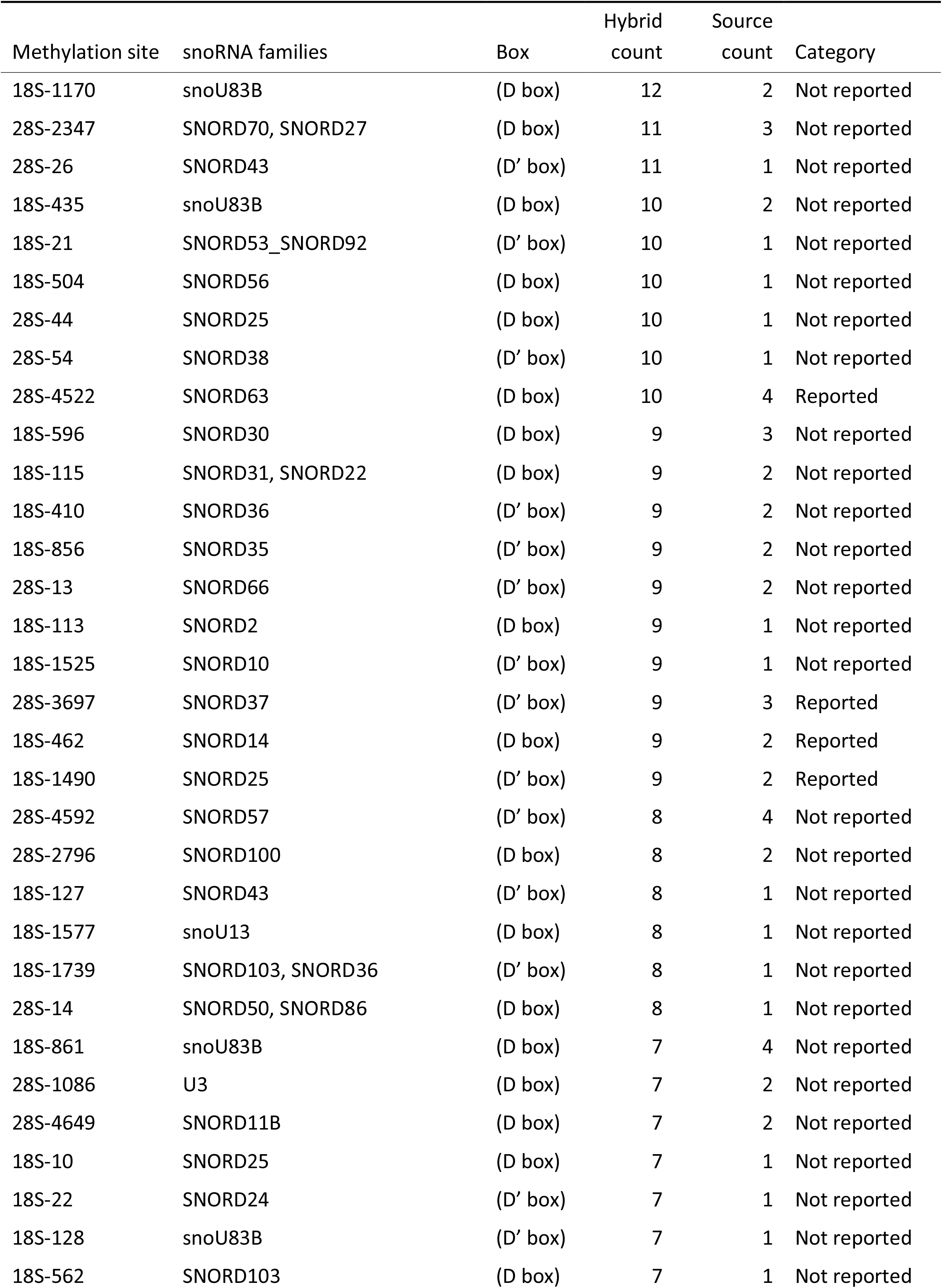
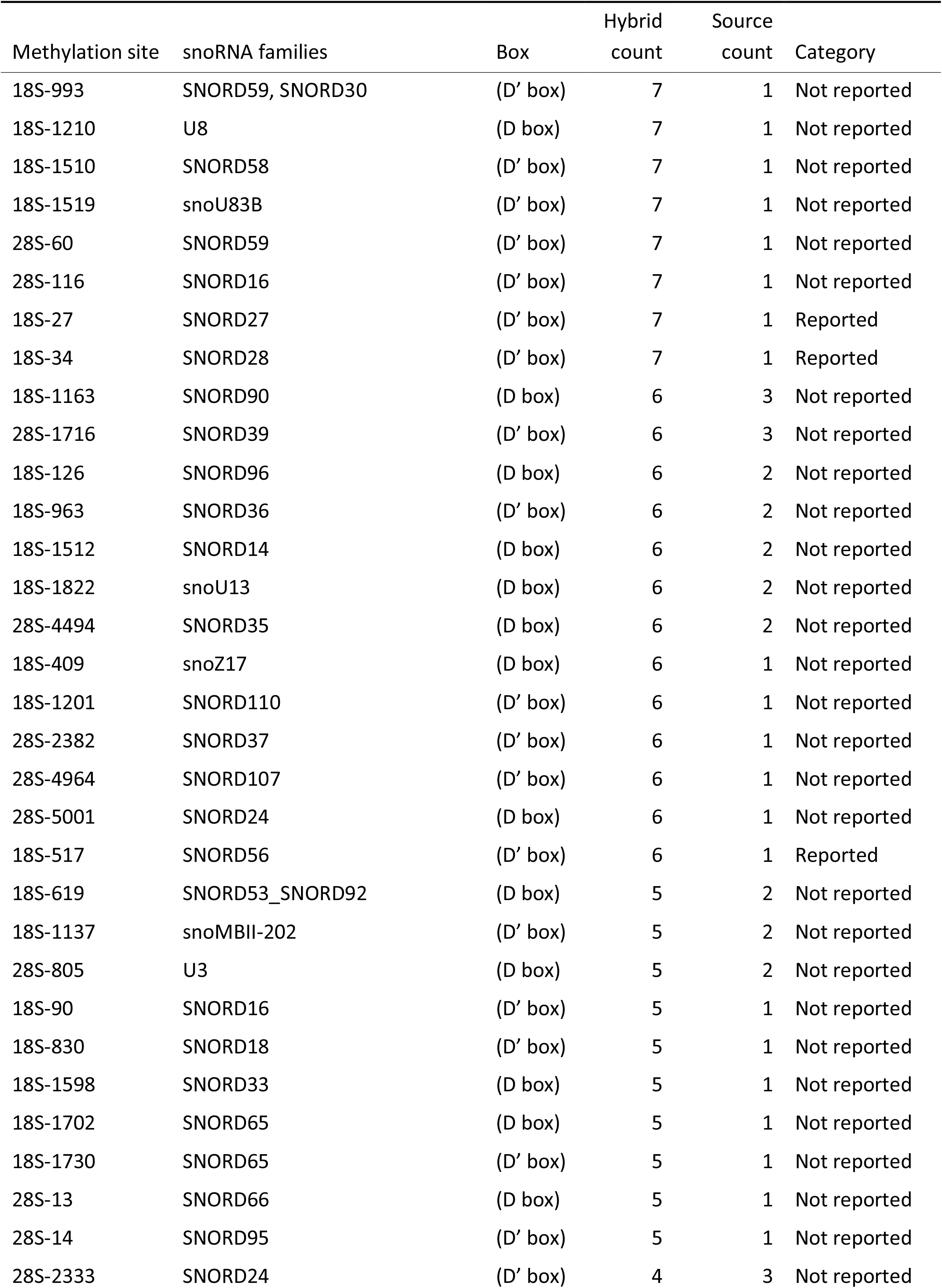
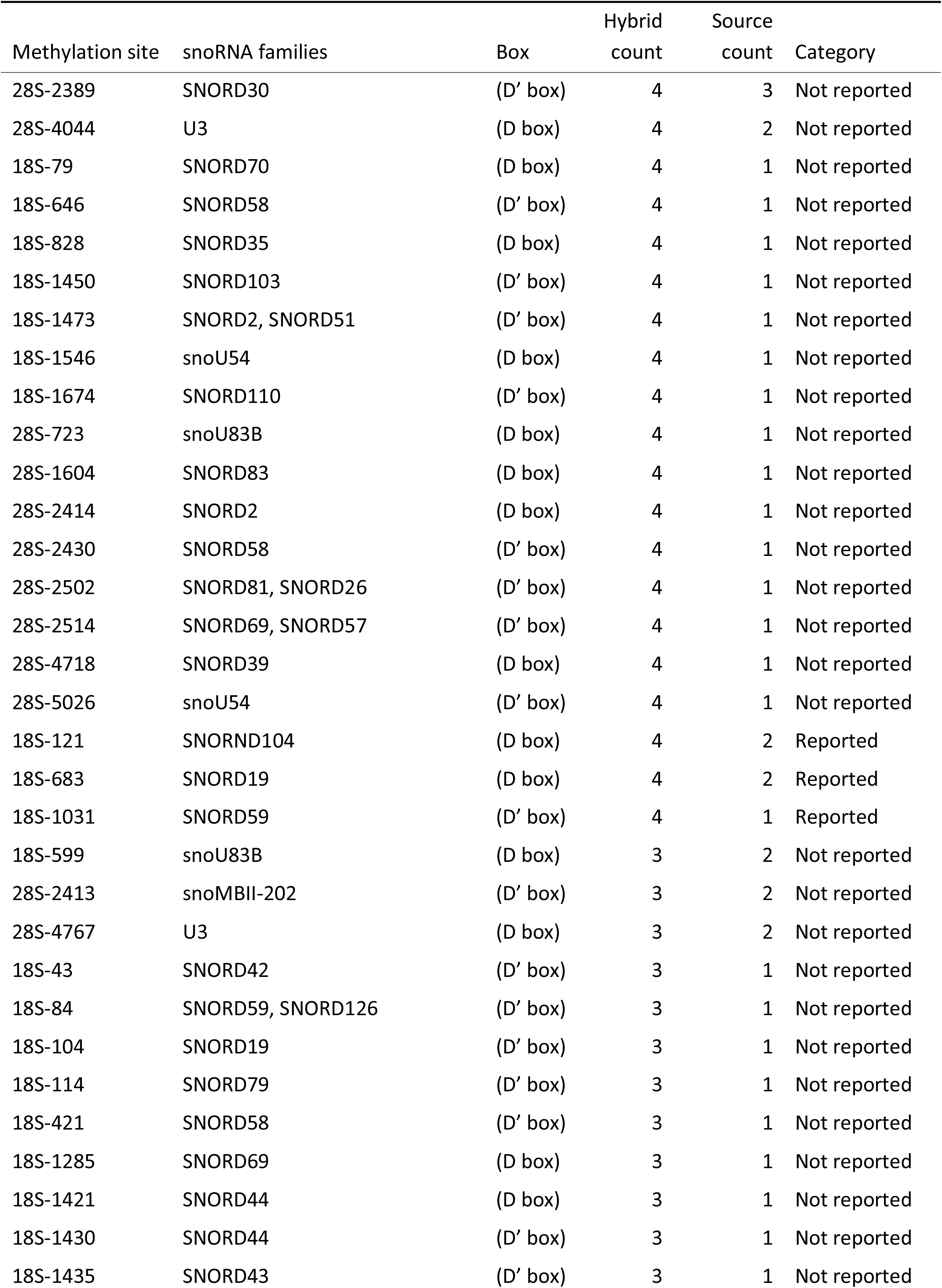
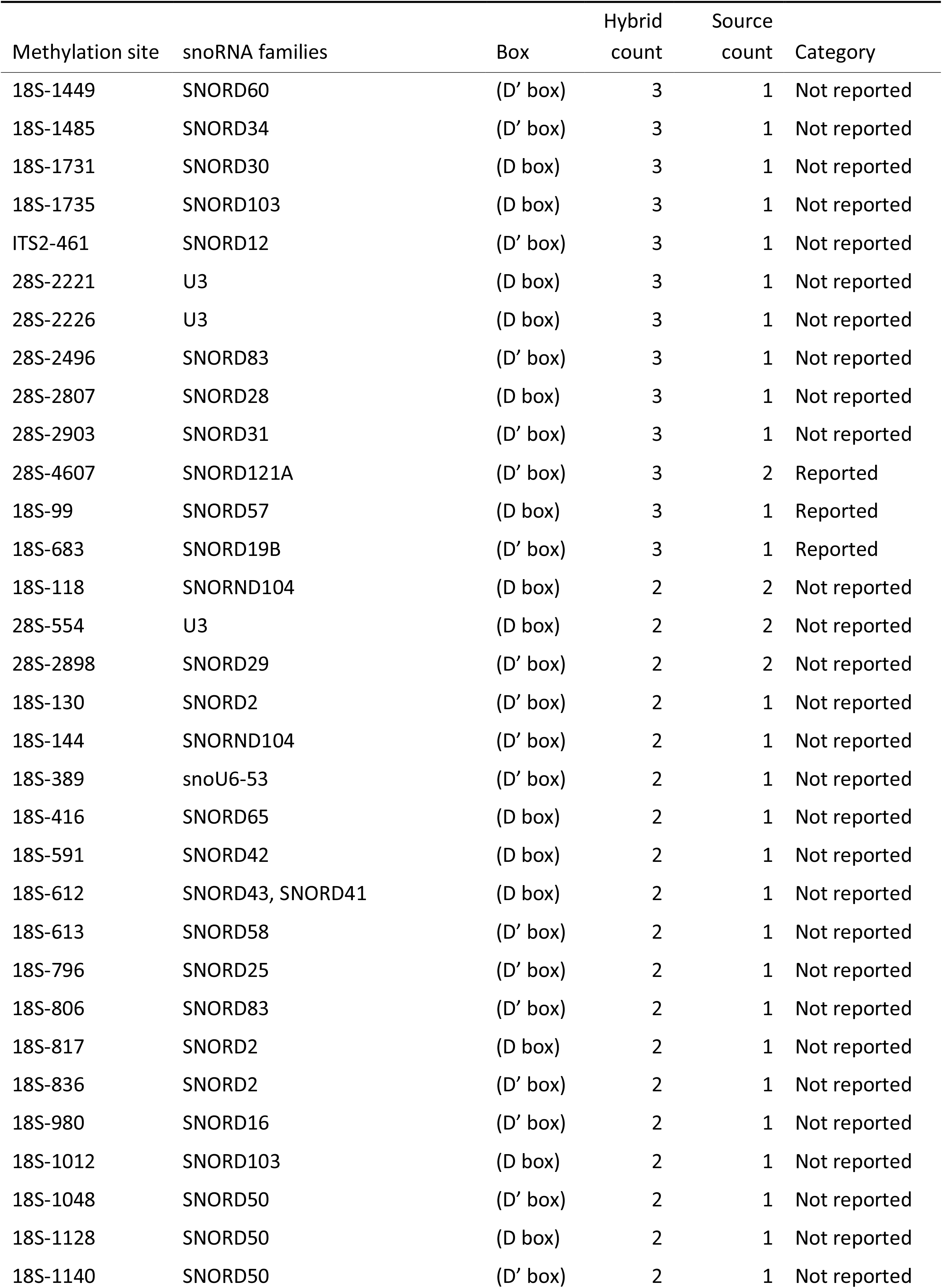
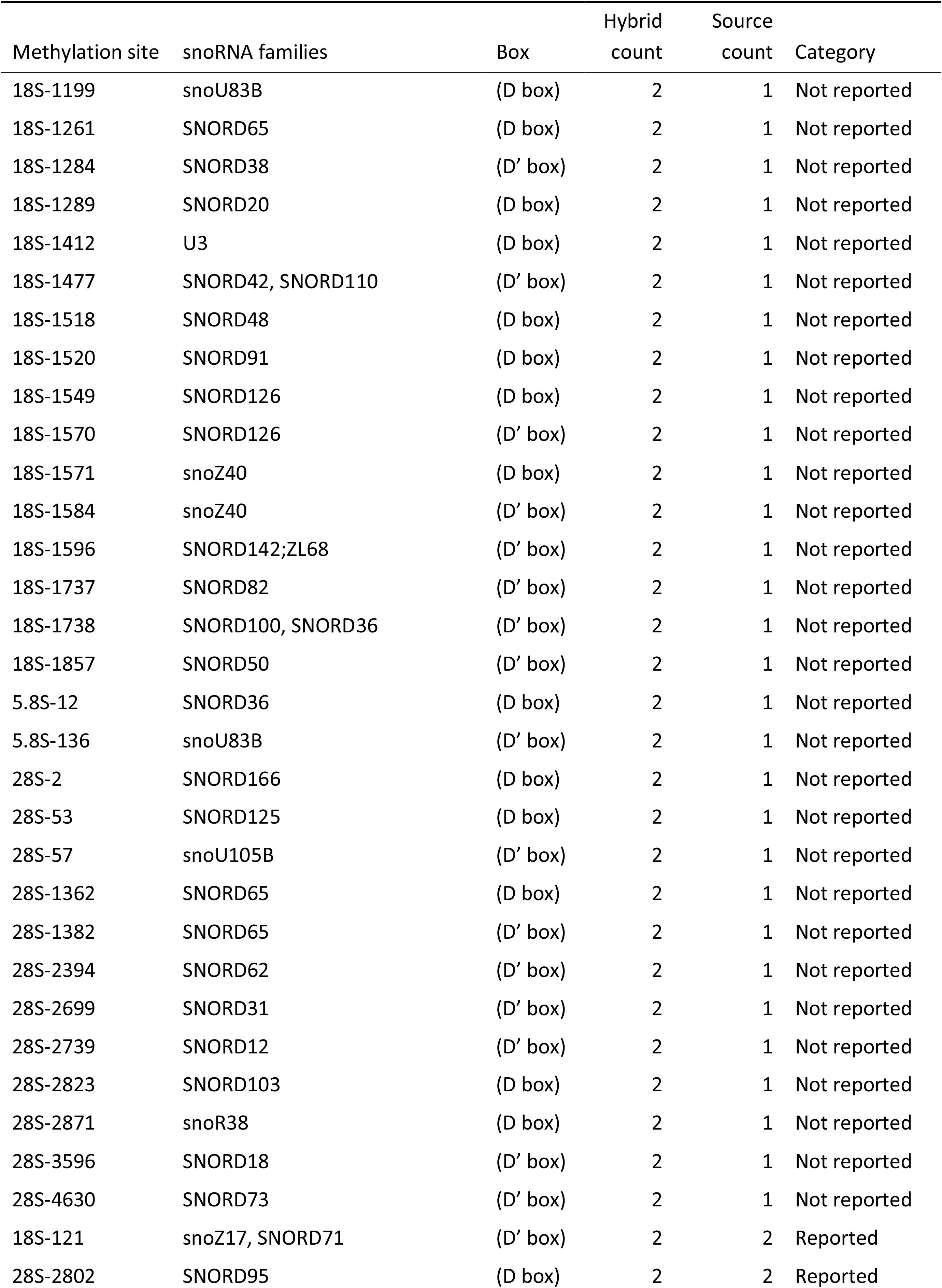
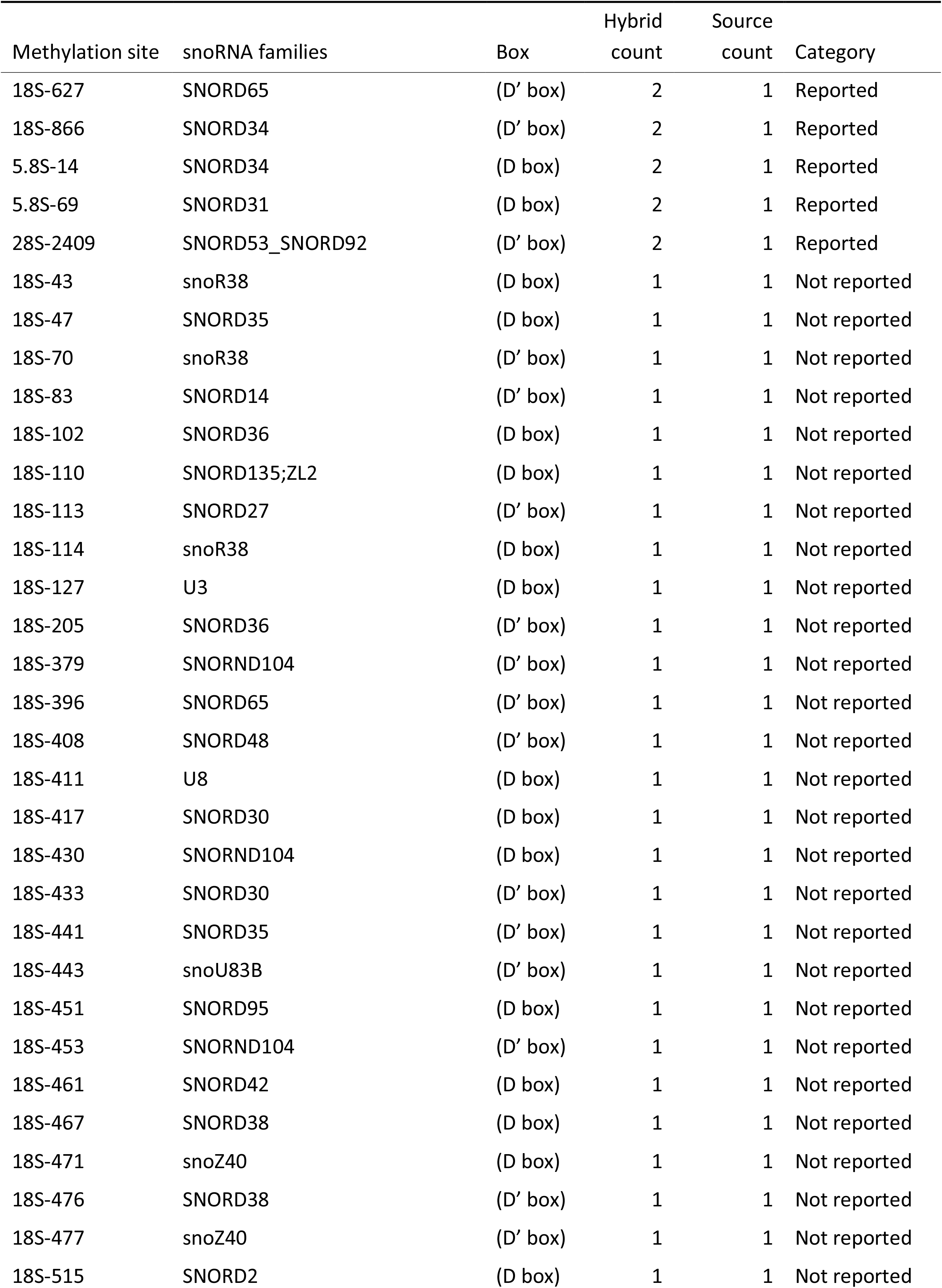
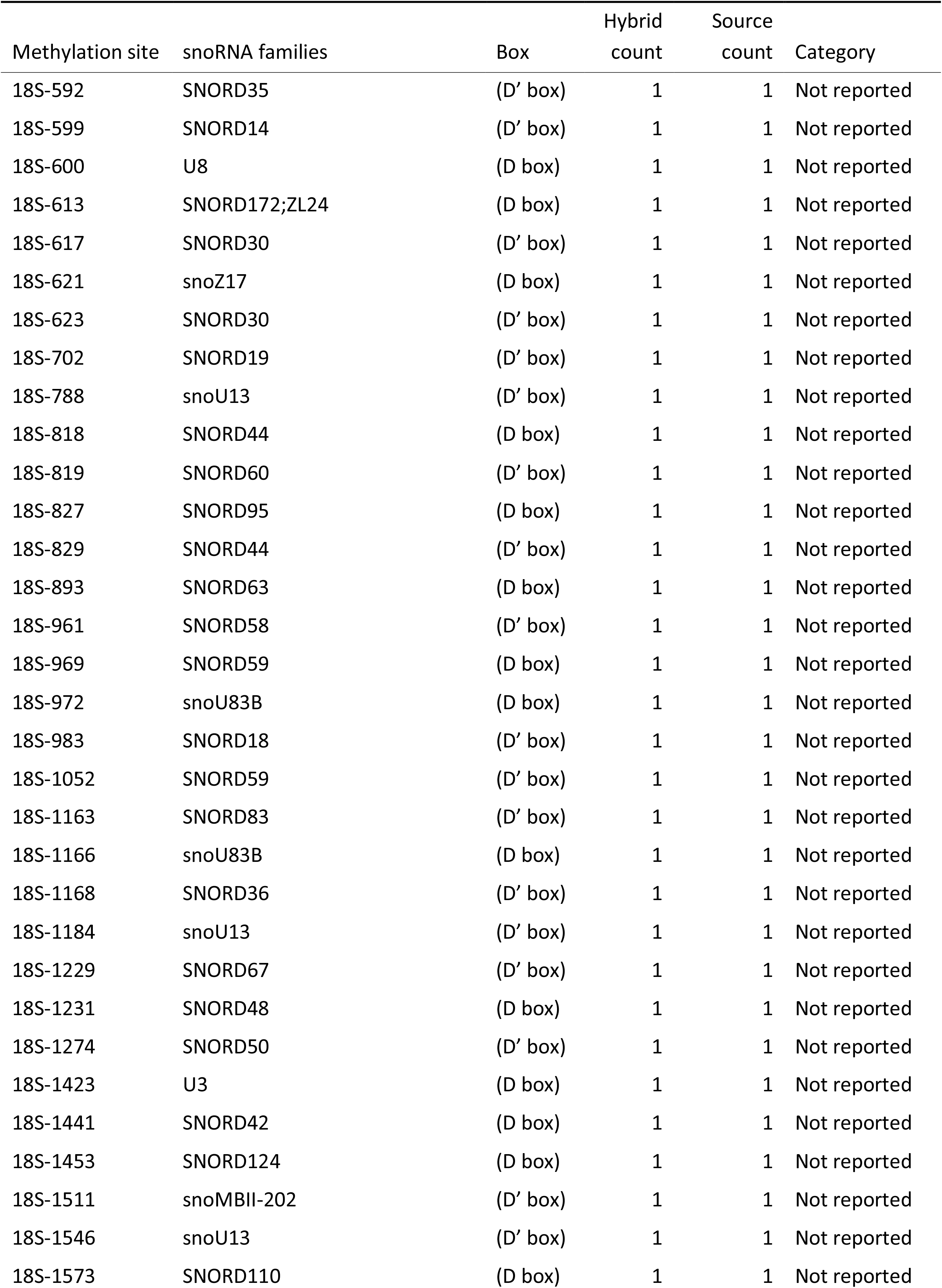
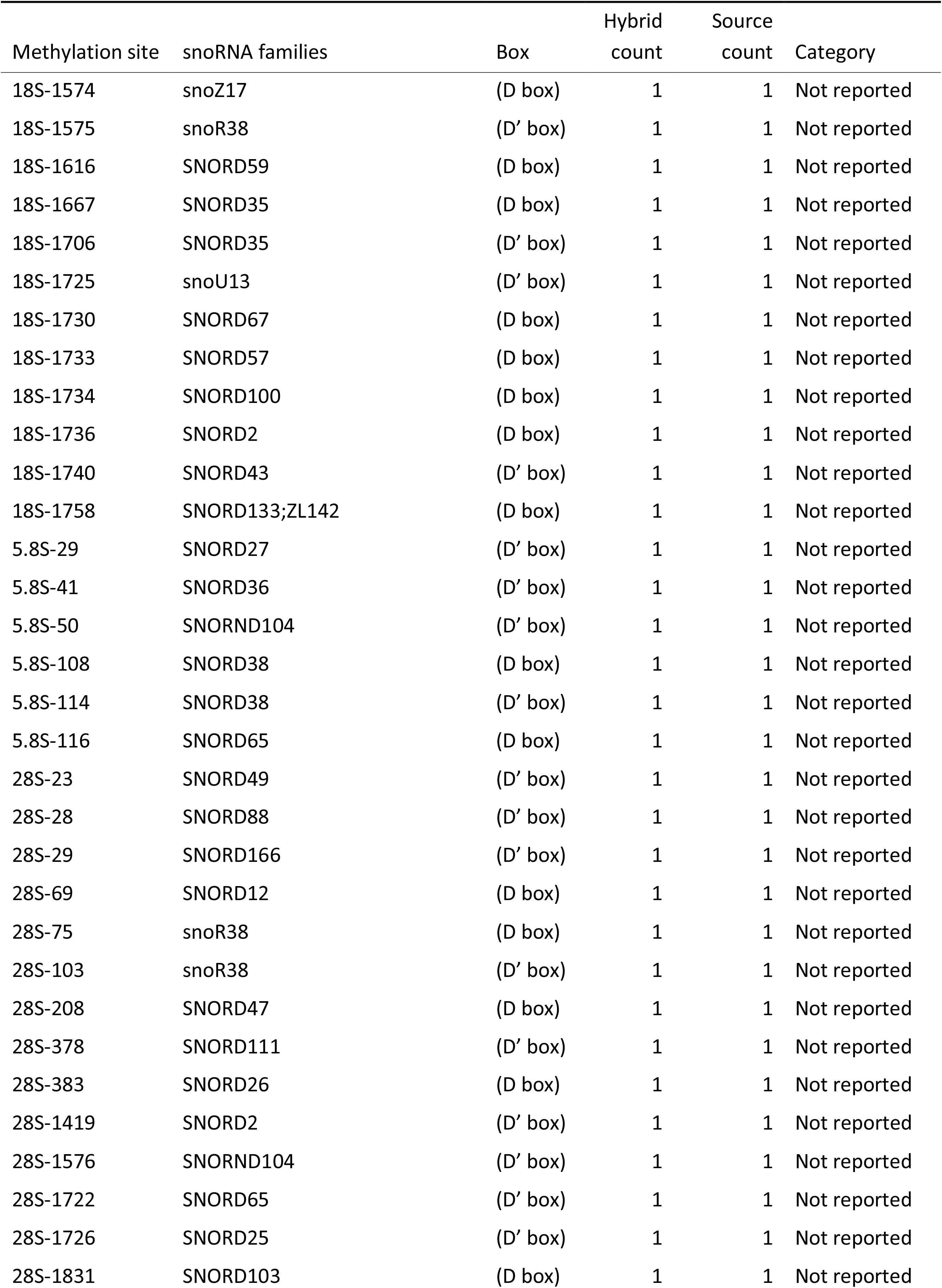
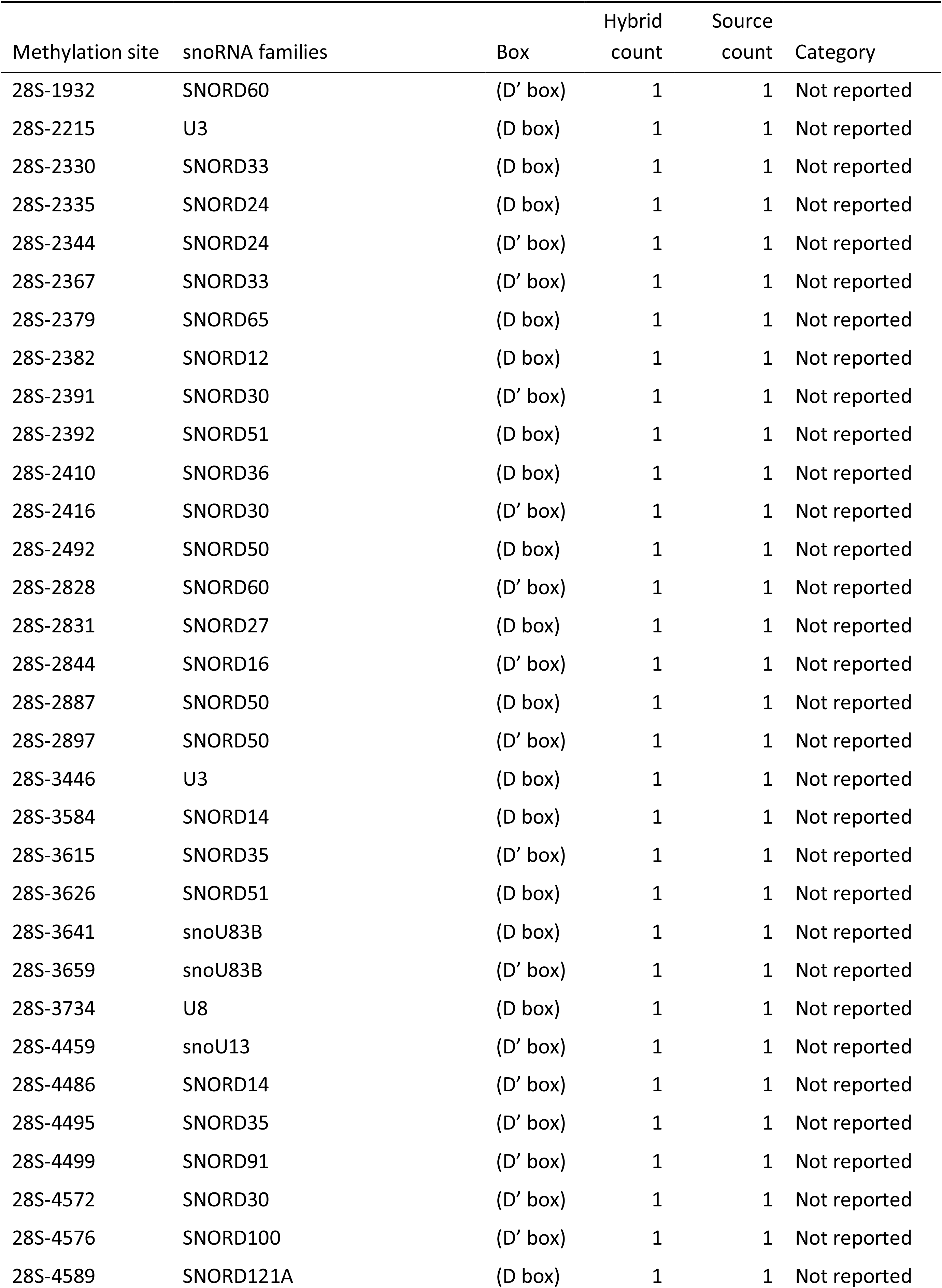
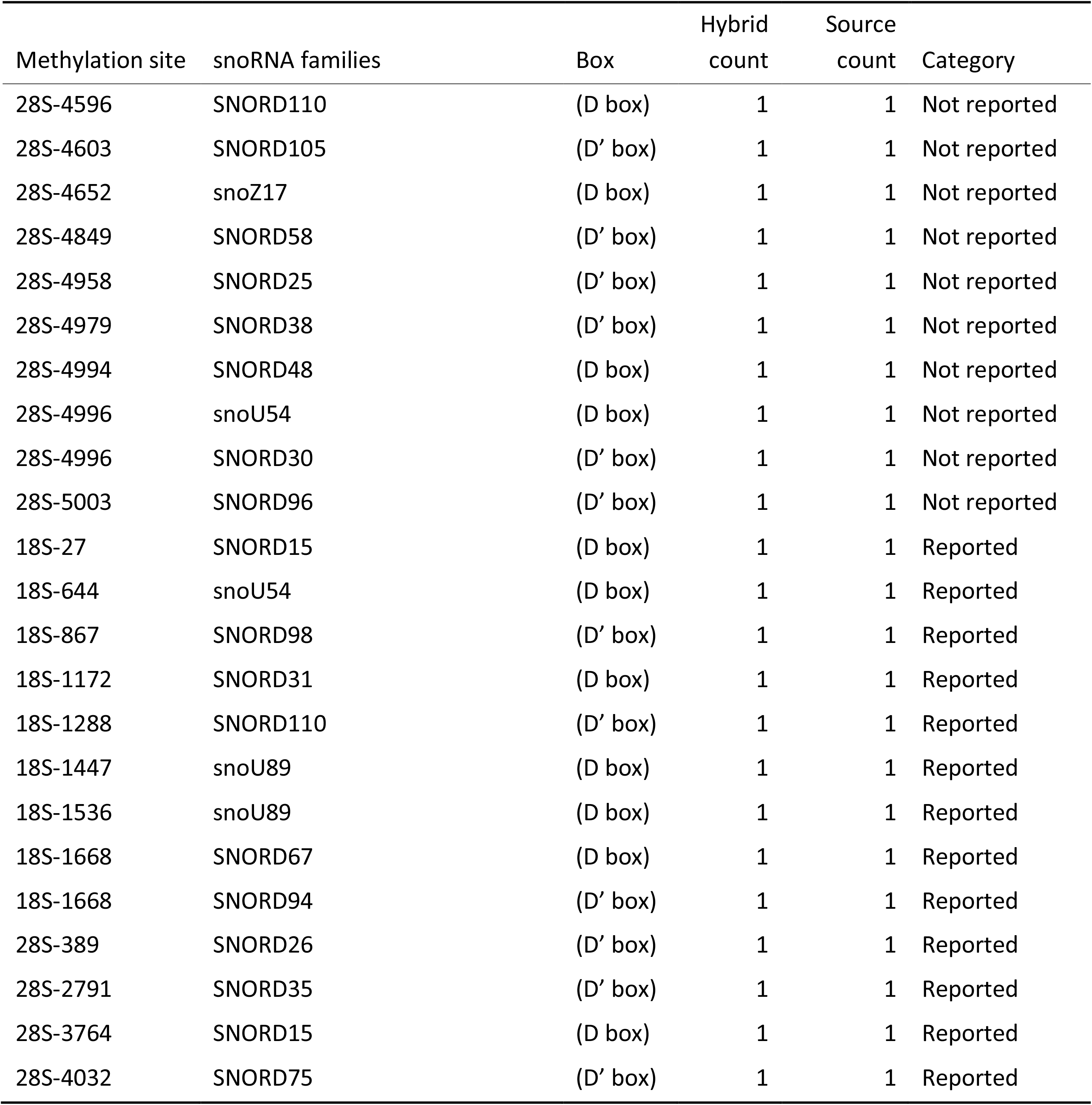
Potentially methylating hybrids.

**Table S4:** Ancillary hybrid counts.

| Interaction | Hybrid count | Experiment count |
| --- | --- | --- |
| SNORD25 : 18S-10 | 35 | 2 |
| SNORD31 : 5.8S-69 | 23 | 1 |
| SNORD48 : 18S-1518 | 19 | 1 |
| SNORD59 : 28S-60 | 19 | 1 |
| SNORD2 : 18S-113 | 17 | 1 |
| SNORD58 : 18S-421 | 17 | 1 |
| SNORD20 : 18S-1804 | 16 | 3 |
| SNORD20 : 18S-1806 | 16 | 3 |
| SNORD35 : 28S-2791 | 16 | 2 |
| SNORD100 : 18S-436 | 13 | 1 |
| SNORD110 : 18S-1477 | 13 | 3 |
| SNORD110 : 18S-1573 | 13 | 3 |
| SNORD126 : 18S-84 | 13 | 1 |
| SNORD2 : 18S-1736 | 13 | 1 |
| snoU83B : 18S-1519 | 13 | 1 |
| SNORD103 : 18S-1668 | 12 | 1 |
| U8 : 18S-1210 | 12 | 1 |
| snoU83B : 28S-3659 | 12 | 1 |
| SNORD25 : 28S-44 | 11 | 1 |
| snoZ17 : 18S-409 | 10 | 2 |
| snoU83B : 28S-3641 | 10 | 1 |
| SNORD35 : 18S-441 | 9 | 1 |
| SNORD35 : 18S-856 | 9 | 4 |
| U3 : 28S-3031 | 9 | 3 |
| SNORD30 : 18S-1731 | 8 | 1 |
| SNORD33 : 18S-1326 | 8 | 3 |
| SNORD24 : 18S-22 | 8 | 1 |
| snoU83B : 18S-1199 | 8 | 1 |
| SNORD103 : 18S-1739 | 8 | 1 |
| SNORD11B : 18S-1605 | 7 | 2 |
| SNORD16 : 28S-2844 | 7 | 1 |
| SNORD35 : 18S-1667 | 7 | 1 |
| SNORD37 : 28S-3697 | 7 | 1 |
| SNORD48 : 18S-1231 | 7 | 2 |
| SNORD65 : 18S-157 | 7 | 1 |
| SNORD2 : 18S-130 | 6 | 1 |
| SNORD31 : 28S-2903 | 6 | 1 |
| SNORD43 : 18S-127 | 6 | 1 |
| SNORD69 : 18S-1285 | 6 | 1 |
| snoU83B : 18S-128 | 6 | 3 |
| SNORD100 : 28S-4576 | 5 | 1 |
| U8 : 18S-411 | 5 | 1 |
| U8 : 18S-412 | 5 | 1 |
| U8 : 18S-1605 | 5 | 1 |
| SNORD14 : 18S-1512 | 5 | 1 |
| SNORD33 : 18S-1328 | 5 | 1 |
| SNORD57 : 18S-99 | 5 | 1 |
| SNORD70 : 18S-79 | 5 | 1 |
| SNORD79 : 18S-114 | 5 | 1 |
| snoU83B : 18S-507 | 5 | 1 |
| snoU83B : 18S-599 | 5 | 1 |
| SNORD103 : 28S-2823 | 5 | 1 |
| SNORD142;ZL68 : 18S-1596 | 4 | 2 |
| SNORD36 : 28S-15 | 4 | 1 |
| SNORD38 : 18S-467 | 4 | 1 |
| SNORD38 : 18S-476 | 4 | 1 |
| SNORD44 : 18S-818 | 4 | 1 |
| SNORD44 : 18S-829 | 4 | 1 |
| snoZ17 : 18S-1574 | 4 | 1 |
| SNORD58 : 18S-1510 | 4 | 1 |
| SNORND104 : 18S-144 | 3 | 1 |
| SNORND104 : 5.8S-50 | 3 | 1 |
| SNORD110 : 18S-1674 | 3 | 2 |
| snoR38 : 18S-70 | 3 | 1 |
| SNORD27 : 28S-2347 | 3 | 1 |
| SNORD30 : 18S-417 | 3 | 1 |
| SNORD30 : 18S-433 | 3 | 1 |
| SNORD36 : 18S-1739 | 3 | 1 |
| SNORD42 : 18S-43 | 3 | 1 |
| SNORD45 : 18S-159 | 3 | 1 |
| SNORD58 : 18S-613 | 3 | 1 |
| SNORD59 : 18S-84 | 3 | 1 |
| snoZ40 : 18S-1571 | 3 | 1 |
| SNORD62 : 28S-2394 | 3 | 2 |
| SNORD63 : 18S-893 | 3 | 1 |
| SNORD66 : 28S-13 | 3 | 2 |
| snoMBII-202 : 18S-1511 | 3 | 1 |
| SNORA46 : 18S-649 | 2 | 1 |
| SNORD100 : 18S-405 | 2 | 1 |
| SNORD100 : 18S-411 | 2 | 1 |
| SNORND104 : 28S-1576 | 2 | 1 |
| snoU105B : 28S-57 | 2 | 1 |
| SNORD12 : 28S-2739 | 2 | 1 |
| snoU13 : 18S-1725 | 2 | 1 |
| SNORD133;ZL142 : 18S-1758 | 2 | 1 |
| SNORD135;ZL2 : 18S-110 | 2 | 1 |
| SNORD15 : 18S-27 | 2 | 1 |
| SNORD19 : 18S-702 | 2 | 1 |
| snoR38 : 28S-2871 | 2 | 1 |
| SNORD2 : 18S-1164 | 2 | 1 |
| SNORD31 : 18S-1172 | 2 | 1 |
| SNORD33 : 18S-1598 | 2 | 1 |
| SNORD34 : 18S-1485 | 2 | 1 |
| SNORD34 : 5.8S-14 | 2 | 1 |
| SNORD35 : 18S-47 | 2 | 2 |
| SNORD35 : 18S-828 | 2 | 1 |
| SNORD36 : 28S-3703 | 2 | 1 |
| SNORD36 : 18S-410 | 2 | 1 |
| SNORD36 : 18S-668 | 2 | 1 |
| SNORD38 : 28S-4979 | 2 | 2 |
| U3 : 28S-3446 | 2 | 2 |
| SNORD42 : 18S-461 | 2 | 1 |
| snoZ17 : 18S-121 | 2 | 1 |
| SNORD39 : 28S-2791 | 2 | 1 |
| SNORD59 : 18S-1616 | 2 | 1 |
| snoZ40 : 18S-1584 | 2 | 1 |
| SNORD65 : 18S-396 | 2 | 1 |
| SNORD65 : 18S-1261 | 2 | 1 |
| SNORD65 : 28S-1722 | 2 | 1 |
| SNORD83 : 18S-806 | 2 | 1 |
| SNORD86 : 18S-1219 | 2 | 2 |
| SNORD100 : 18S-1734 | 1 | 1 |
| SNORD100 : 18S-1738 | 1 | 1 |
| U8 : 18S-600 | 1 | 1 |
| SNORD12 : 18S-1536 | 1 | 1 |
| SNORD14 : 18S-462 | 1 | 1 |
| snoR38 : 18S-114 | 1 | 1 |
| snoR38 : 18S-1575 | 1 | 1 |
| snoR38 : 18S-43 | 1 | 1 |
| SNORD25 : 18S-1490 | 1 | 1 |
| SNORD30 : 18S-1383 | 1 | 1 |
| SNORD36 : 18S-205 | 1 | 1 |
| SNORD36 : 18S-1738 | 1 | 1 |
| SNORD45 : 18S-159 | 1 | 1 |
| SNORD65 : 18S-416 | 1 | 1 |
| snoMBII-202 : 18S-428 | 1 | 1 |
| SNORD73 : 28S-1747 | 1 | 1 |
| snoU83B : 18S-1199 | 1 | 1 |
| SNORD91 : 28S-4588 | 1 | 1 |
| SNORD53_SNORD92 : 18S-619 | 1 | 1 |

**Table S5:** Blocking hybrid counts.

| Interaction | Hybrid count | Experiment count |
| --- | --- | --- |
| snoZ17 : 28S-4589 | 9 | 1 |
| snoZ17 : 28S-4588 | 8 | 1 |
| SNORD91 : 18S-1052 | 8 | 2 |
| snoZ17 : 28S-4592 | 4 | 1 |
| snoZ17 : 28S-4593 | 4 | 1 |
| SCARNA17 : 18S-93 | 3 | 1 |
| SNORD33 : 18S-93 | 1 | 1 |
| SNORD91 : 18S-1056 | 4 | 1 |
| snoZ17 : 28S-4596 | 3 | 1 |
| snoZ17 : 28S-4598 | 3 | 1 |
| SCARNA17 : 18S-95 | 2 | 1 |
| SNORD33 : 18S-95 | 1 | 1 |
| SNORD34 : 18S-22 | 2 | 1 |
| SNORD34 : 18S-27 | 2 | 1 |
| SNORD34 : 18S-28 | 2 | 1 |
| SCARNA17 : 18S-90 | 1 | 1 |
| SNORD33 : 18S-90 | 1 | 1 |
| snoZ17 : 28S-4590 | 1 | 1 |
| snoZ17 : 28S-4603 | 1 | 1 |
| SNORD34 : 18S-21 | 1 | 1 |
| SNORD33 : 18S-84 | 1 | 1 |

**Table S6:** Structural hybrid counts.

| Interaction | Hybrid count | Experiment count |
| --- | --- | --- |
| U3 | 6,827 | 14 |
| U8 | 2,118 | 12 |
| snoU83B | 1,124 | 11 |
| SNORD58 | 1,022 | 3 |
| SNORD14 | 834 | 11 |
| snoU13 | 812 | 13 |
| SNORD62 | 580 | 7 |
| SNORD38 | 543 | 6 |
| SNORD91 | 501 | 4 |
| snoR38 | 472 | 4 |
| SNORD103 | 429 | 4 |
| SNORD44 | 419 | 1 |
| SNORD83 | 416 | 5 |
| SNORD107 | 396 | 2 |
| SNORD100 | 394 | 7 |
| SNORD36 | 380 | 3 |
| SNORD110 | 342 | 7 |
| SNORD34 | 321 | 3 |
| SNORD2 | 310 | 2 |
| snoZ17 | 307 | 3 |
| SNORD16 | 301 | 6 |
| SNORD45 | 295 | 3 |
| snoMBII-202 | 294 | 6 |
| SNORD31 | 255 | 5 |
| snoU2_19 | 251 | 5 |
| others | 8,057 | 16 |

